# A phospho-regulated signal motif determines subcellular localization of α-TAT1 for dynamic microtubule acetylation

**DOI:** 10.1101/2020.09.23.310235

**Authors:** Abhijit Deb Roy, Evan G. Gross, Gayatri S. Pillai, Shailaja Seetharaman, Sandrine Etienne-Manneville, Takanari Inoue

**Affiliations:** Department of Cell Biology and Center for Cell Dynamics, Johns Hopkins University School of Medicine, 855 North Wolfe Street, Baltimore, MD 21205, USA; The Johns Hopkins University, Baltimore, MD 21218, USA; Cell Polarity, Migration and Cancer Unit, Institut Pasteur, UMR3691 CNRS, Equipe Labellisée Ligue Contre le Cancer, F-75015, Paris, France; Université Paris Descartes, Sorbonne Paris Cité, 12 Rue de l’École de Médecine, 75006 Paris, France

## Abstract

Spatiotemporally dynamic microtubule acetylation underlies diverse physiological events ranging from cell migration to intracellular trafficking, autophagy and viral infections. Despite its ubiquity, the molecular mechanisms that regulate the sole microtubule acetylating agent, α-tubulin N-acetyltransferase 1 (α-TAT1) remain obscure. Here we report that dynamic intracellular localization of α-TAT1 unexpectedly determines the efficiency of microtubule acetylation. Specifically, we newly identified a conserved signal motif in the intrinsically disordered C-terminus of α-TAT1, consisting of three competing regulatory elements - nuclear export, nuclear import and cytosolic retention. Their balance is tuned via phosphorylation by serine-threonine kinases including CDK1 and CK2. While the un-phosphorylated form resides both in the cytosol and nucleus, the phosphorylated form binds to specific 14-3-3 adapters and accumulates in the cytosol for maximal substrate access. Cytosolic localization of α-TAT1 predominantly mediates microtubule acetylation, cell proliferation and DNA damage response. In contrast to other molecules with a similar phospho-regulated signal motif including transcription factors, α-TAT1 uniquely uses the nucleus as a hideout. As amino acid mutations to the motif have been reported in cancer patients, the present mechanism of subcellular α-TAT1 localization may help uncover a spatiotemporal code of microtubule acetylation in normal and aberrant cell functions.

## Introduction

Acetylation of Lysine-40 of α-tubulin is an evolutionarily conserved post-translational modification observed across eukaryotic species^1–3^, which is involved in diverse physiological and pathological states^4–6^. Acetylation is mainly observed in polymerized microtubules^7–9^ and may provide structural flexibility to stabilize microtubules against bending forces^10–13^. In cultured cells, microtubule acetylation mediates focal adhesion dynamics, adaptation to extracellular matrix rigidity as well as regulation of tissue stiffness^14–17^. Additionally, acetylated microtubules regulate touch sensation in *M. musculus, C. elegans* and *D. melanogaster*, suggesting a role in mechano-response^18–21^. Microtubule acetylation has been implicated in axonal transport in neurons^22–24^, migration in cancer cells^5,25–27^, autophagy^6,28,29^, podosome stabilization in osteoclasts^30^ and viral infections^31–34^. α-TAT1 is the only known acetyltransferase for α-tubulin^35,36^ in mammals. α-TAT1 predominantly catalyzes α-tubulin in stable polymerized microtubules^19,37^, and may have additional effects on microtubules independent of its catalytic activity^38^.

Although microtubule acetylation is spatially and temporally regulated downstream of many molecular signaling pathways, little is known about how these pathways converge on α-TAT1 to achieve such dynamic patterns. In the present study, we used computational sequence analyses and live cell microscopy to identify a conserved motif in the intrinsically disordered C-terminus of α-TAT1, consisting of an NES and an NLS, that mediates its spatial distribution. We show that cytosolic localization of α-TAT1 is critical for microtubule acetylation. We further demonstrate that nuclear localization of α-TAT1 is inhibited by the action of serine-threonine kinases, specifically cyclin dependent kinases (CDKs), protein kinase A (PKA) and casein kinase 2 (CK2) and identify 14-3-3 proteins as binding partners of α-TAT1. Our findings establish a novel role of the intrinsically disordered C-terminus in controlling α-TAT1 function by regulating its intracellular localization downstream of kinase and phosphatase activities.

## Results

### α-TAT1 localization mediates microtubule acetylation

α-TAT1 has an N-terminal catalytic domain that shows homology to other acetyltransferases, while its C-terminus was not resolved in crystal structures^39^ (Fig. 1a). Based on its amino acid sequence, α-TAT1 C-terminus was predicted to be intrinsically disordered by both IUPred2A^40^ and PrDOS^41^ prediction servers (Fig. 1a, Supplementary Fig. S1a, b, c). To explore if the intra-cellular localization of α-TAT1 is regulated, we sought to identify any localization signals present in the α-TAT1 amino acid sequence (Supplementary Fig. S1a). The prediction program NetNES^42^ identified a putative NES in α-TAT1 C-terminus (Fig. 1a, Supplementary Fig. S2). On the other hand, PSORT-II^43^ subcellular localization program predicted that α-TAT1 should be predominantly localized to the nucleus due to the presence of a putative class 4 NLS^44^ in its C-terminus (Fig. 1a). The region encompassing the putative NES and NLS is conserved across the human α-TAT1 isoforms (Supplementary Fig. S3a), as well as across mammalian α-TAT1 proteins (Supplementary Fig. S3b). To test whether α-TAT1 indeed showed any intracellular distribution pattern, we expressed mVenus-α-TAT1 in HeLa cells, and observed distinct nuclear exclusion in most cells, although a subset of cells showed lack of exclusion (Fig. 1b). Based on visual categorization (see Methods, Supplementary Fig.S4a), we determined that approximately 80% of cells showed cytosolic distribution and 20% showed lack of nuclear exclusion (diffused pattern or nuclear enrichment) of mVenus-α-TAT1 (Fig. 1c). These observations were consistent with ratiometric and co-localization correlation analyses (see Methods, Supplementary Fig. S4a) of mVenus-α-TAT1 distribution (Fig. 1d, e). The nuclear/cytosolic ratio did not show a strong correlation with mVenus-α-TAT1 expression levels (Supplementary Fig. S4b), suggesting that the intracellular localization was not an artifact of exogenous expression of mVenus-α-TAT1. Time-lapse microscopy showed temporal changes in fluorescence intensity of mVenus-α-TAT1 in cell nuclei and cytosol (Supplementary Fig. S5a, b), suggesting that α-TAT1 localization is dynamic.

**Figure 1.**
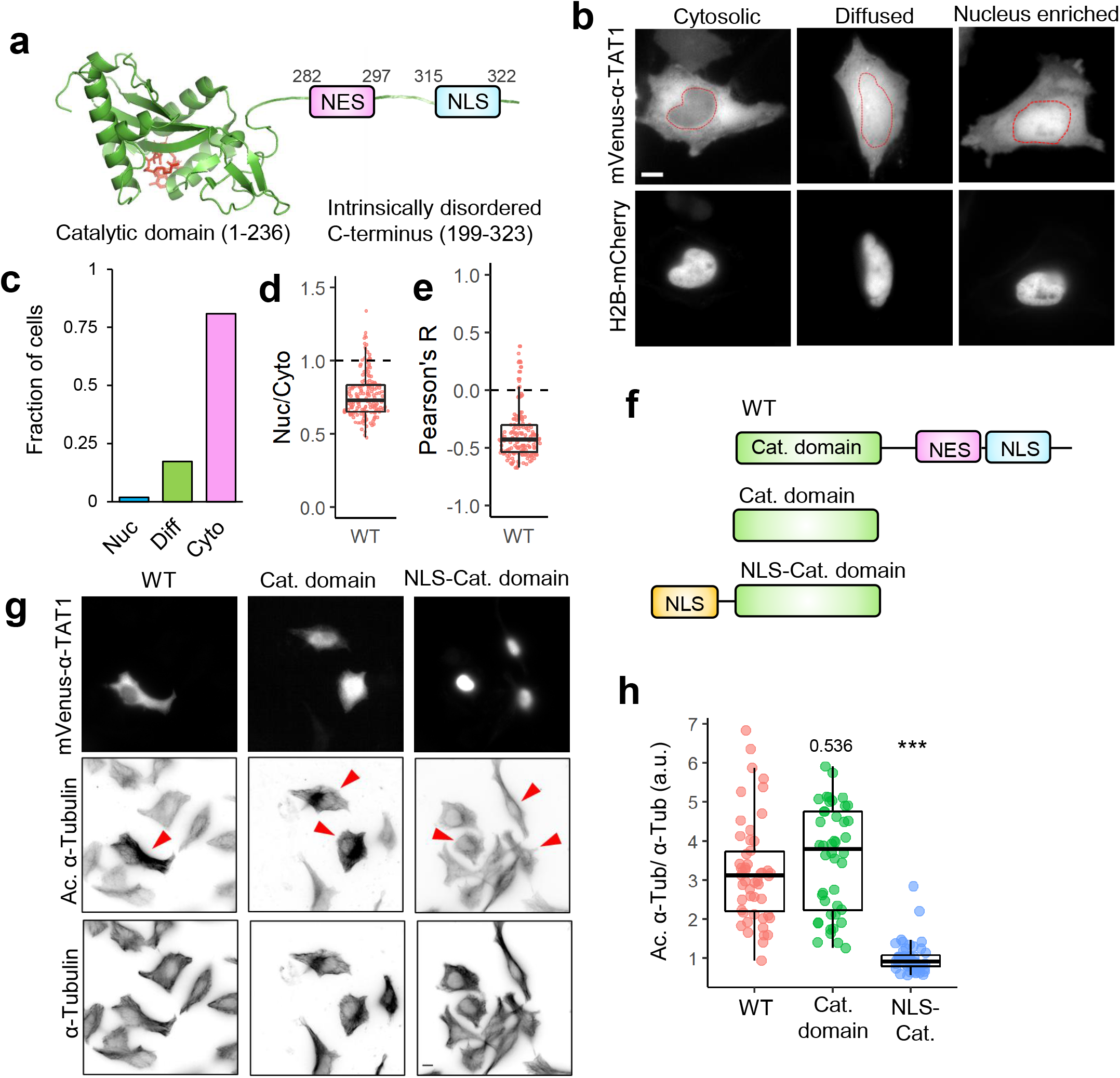
Intracellular distribution of α-TAT1 mediates its function. a) Cartoon showing predicted NES and NLS in intrinsically disordered C-terminus of α-TAT1, adapted from PDB: 4GS4, b) intracellular distribution of mVenus-α-TAT1, red dotted lines outline nuclei as identified by H2B-mCherry in lower panel, c) categorical analysis (318 cells), d) ratiometric analysis (184 cells) and e) colocalization analysis (180 cells) of mVenus-α-TAT1 localization, f) cartoon showing α-TAT1 mutants used in g) immunofluorescence assays showing levels of acetylated α-tubulin and total α-tubulin, transfected cells are indicated with red arrowheads, h) ratio of acetylated α-tubulin to total tubulin intensities with exogenous expression of α-TAT1 and its mutants, normalized against that of non-transfected cells, (WT: 50, catalytic domain: 44, NLS-catalytic domain: 48 cells). Scale bar = 10μm. P-value: *** <0.001 or as shown, Student’s *t*-test.

α-TAT1 preferentially acetylates polymerized microtubules, which are typically cytosolic. Based on this, we hypothesized that spatial regulation of α-TAT1 may control its function. Exogenous expression of mVenus-α-TAT1 or its catalytic domain (residues 1-236)^19,39^ (Fig. 1f) was sufficient to significantly increase α-tubulin acetylation in HeLa cells compared to non-transfected cells (Fig. 1g, h). To test whether nuclear localization may sufficiently sequester α-TAT1 from microtubules, we tethered the NLS from cMyc to α-TAT1 catalytic domain and thus localized it to the nucleus (Fig. 1f, g top panel). Exogenous expression of NLS-mVenus-α-TAT1(1-236) did not increase α-tubulin acetylation levels compared to non-transfected cells (Fig. 1 g, h), suggesting that nuclear sequestration of α-TAT1 inhibits its function.

### α-TAT1 undergoes Exportin 1 dependent nuclear export

To dissect the molecular mechanisms of α-TAT1 localization in the cytosol, we speculated that α-TAT1 is actively exported out of the nucleus, and/or that α-TAT1 binds to a protein that keeps the complex out of the nucleus. We began testing the first possibility by assessing involvement of the nuclear export machinery. Exportin 1 (Exp1), also called Chromosome region maintenance 1 protein homolog (CRM1), mediates nuclear export of many proteins^45,46^. GFP-α-TAT1, but not GFP, co-immunoprecipitated with endogenous Exp1 (Fig. 2a), indicating an interaction between these two proteins. Treatment with 100 nM Leptomycin-B (LMB), an inhibitor of Exp1 mediated nuclear export^47^, significantly decreased the number of cells displaying nuclear exclusion of mVenus-α-TAT1 compared to vehicle (Fig 2b-e). Inhibition of nuclear export was initiated within an hour of LMB treatment, although some cells were refractory to the treatment (Supplementary Fig. S6a, b). Decreased LMB concentrations (1 nM and 10 nM) had a comparable impact as 100 nM dosage (Supplementary Fig. S6c, d). Furthermore, LMB treatment induced significant reduction in α-tubulin acetylation levels in HeLa cells within 4 hours compared to vehicle (Fig. 2f, g). Our data suggest that α-TAT1 is actively exported from the nucleus to the cytosol in an Exp1 dependent manner and that this export facilitates α-TAT1 function. The residual exclusion of α-TAT1 could be due to Exp1-independent nuclear export pathways or association with other cytosolic proteins, which was tested later (see below).

**Figure 2.**
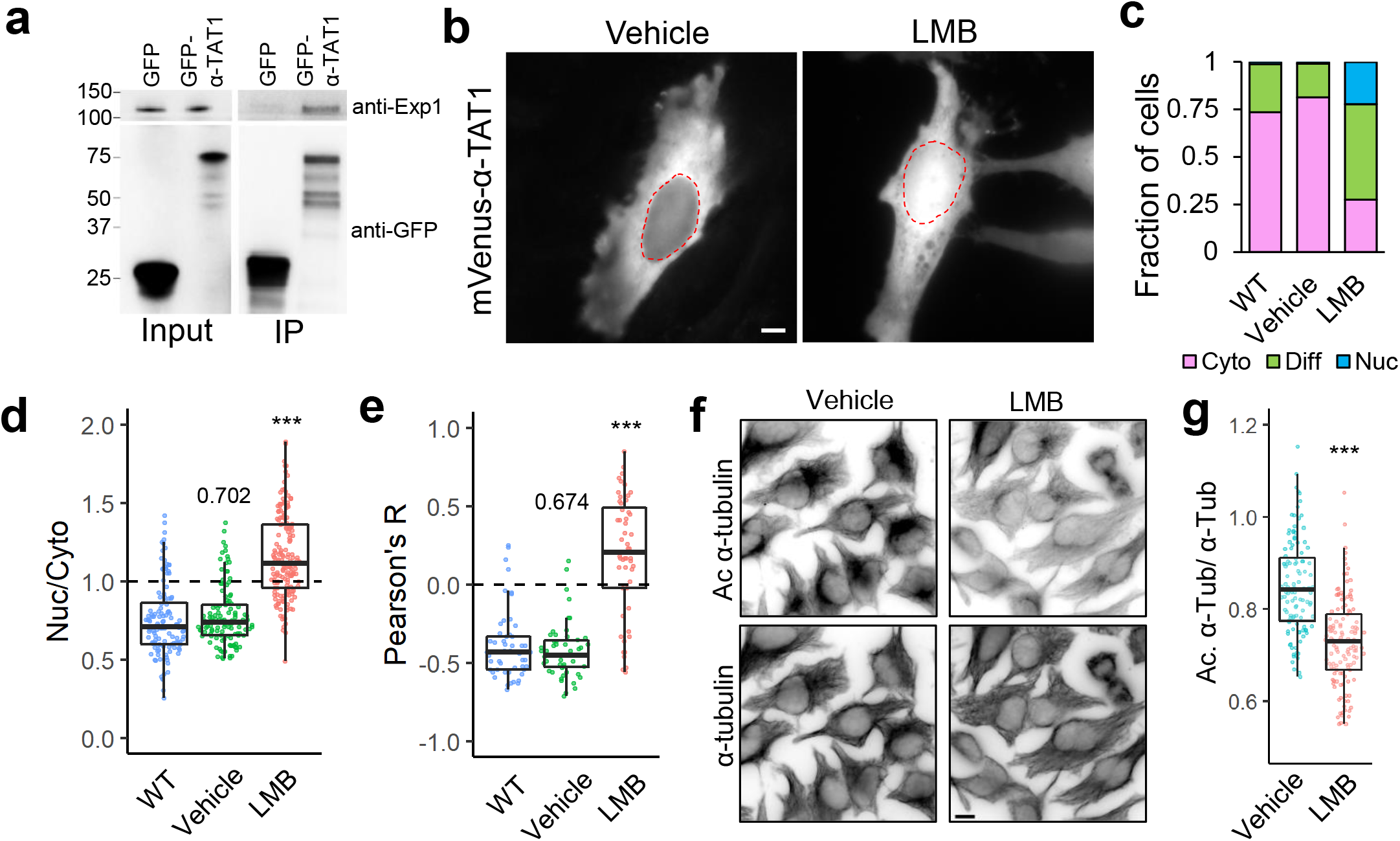
α-TAT1 undergoes Exp1 mediated nuclear export. a) Co-immunoprecipitation of endogenous Exp1 with GFP-α-TAT1, b) intracellular distribution of mVenus-α-TAT1 with vehicle (EtOH) and 100 nM LMB treatment, nuclei are indicated in red dotted lines, c) categorical analysis (WT: 263, vehicle: 300, LMB: 255 cells), d) ratiometric analysis (WT: 125, vehicle: 116, LMB: 163 cells) and e) colocalization analysis (WT: 55, vehicle: 54, LMB: 52 cells) of mVenus-α-TAT1 localization with vehicle and LMB treatment, f) immunofluorescence images showing acetylated and total α-tubulin in HeLa cells with vehicle or LMB treatment, g) ratio of acetylated to total α-tubulin with vehicle or LMB treatment (vehicle:120, LMB: 130 cells). Scale bar = 10μm. P-value: *** <0.001 or as shown, Student’s *t*-test.

### Nuclear export of α-TAT1 is mediated by a C-terminal NES

Our data demonstrate that α-TAT1 function is linked to Exp1 mediated nuclear export. To examine whether nuclear export of α-TAT1 is regulated by its catalytic activity, we expressed a catalytic dead mutant, mVenus-α-TAT1(D157N)^39^ or the catalytic domain, mVenus-α-TAT1(1-236), in HeLa cells. mVenus-α-TAT1(D157N) did not display any loss of nuclear exclusion, whereas mVenus-α-TAT1(1-236), displayed a complete loss of nuclear exclusion (Fig. 3a-e). On the other hand, the C-terminus, mVenus-α-TAT1(236-323), displayed a distribution pattern comparable to that of WT (Fig. 3a-e). In addition, inhibition of Exp1 by 100 nM LMB significantly reduced the nuclear exclusion of mVenus-α-TAT1 C-terminus (Fig. 3f, g). Exclusion of α-TAT1 C-terminus (size ≈ 38 kDa) further indicates that nuclear exclusion of mVenus-α-TAT1 (size ≈ 63 kDa) is not simply due to size exclusion of passive diffusion into nuclei and demonstrates that nuclear exclusion of α-TAT1 is a transferable property mediated by its C-terminus.

**Figure 3.**
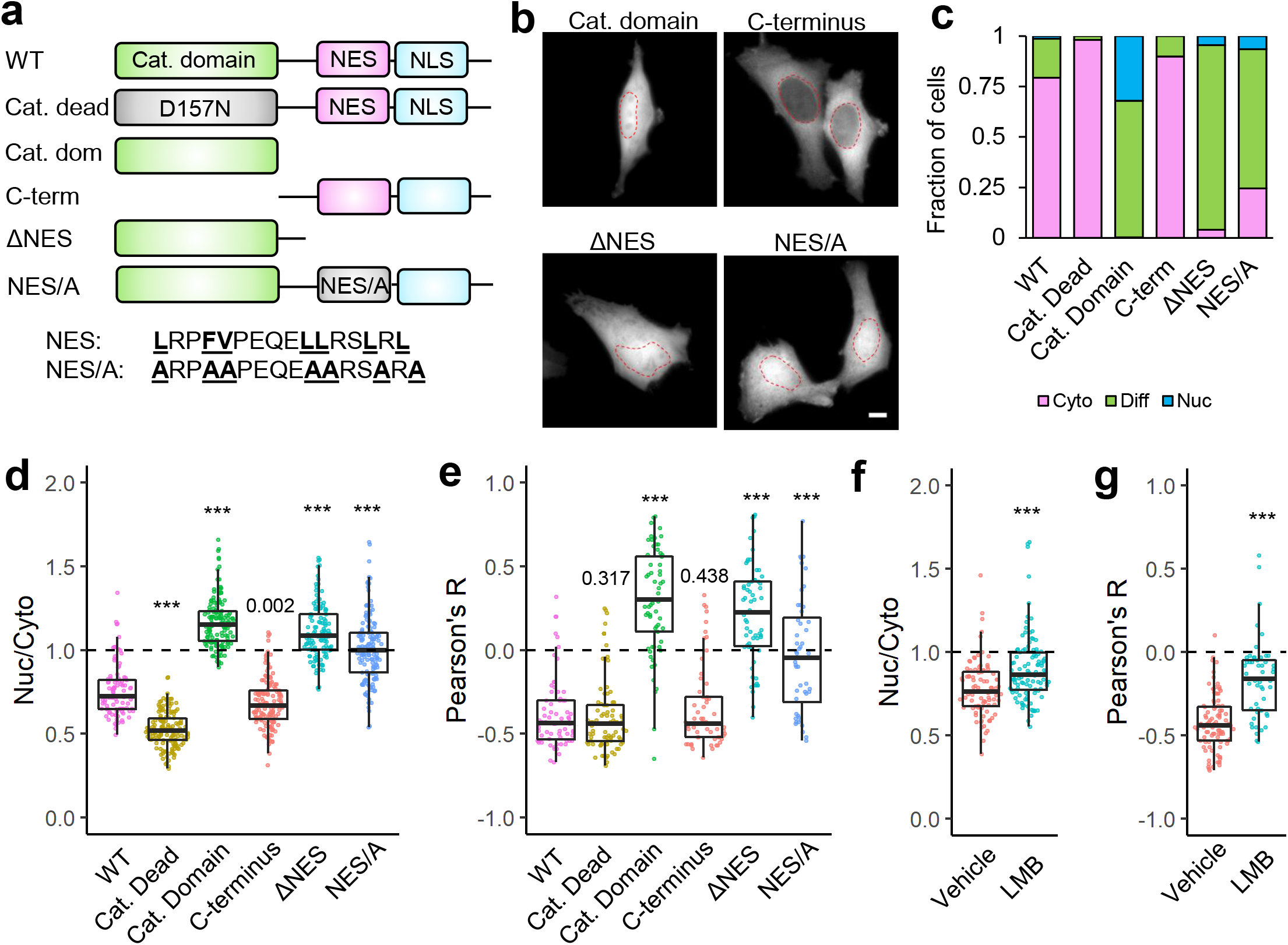
Intracellular distribution of α-TAT1 is mediated by its C-terminal NES. a) Cartoon showing α-TAT1 mutant design, b) representative images showing intracellular distribution of mVenus-α-TAT1 mutants as listed, red dotted lines outline nuclei, c) categorical analysis (WT: 257, Cat. Dead: 289, Cat. Dom: 271, C-term: 290, δNES: 243 and NES/A: 321 cells), d) ratiometric analysis (WT: 103, Cat. Dead: 153, Cat. Dom: 145, C-term: 138, δNES: 124 and NES/A: 153 cells) and e) colocalization analysis (WT: 68, Cat. Dead: 85, Cat. Dom: 65, C-term: 60, δNES: 62 and NES/A: 48 cells) of mVenus-α-TAT1 mutant localization as indicated, f) ratiometric analysis (vehicle: 93, LMB: 107 cells) and g) colocalization analysis (vehicle: 98, LMB: 53) of intracellular distribution of mVenus-α-TAT1 C-term with 100 nM LMB. Scale bar = 10μm. P-value: *** <0.001 or as shown, Student’s *t*-test.

Exp1 dependent nuclear export is typically mediated by binding with short stretches of hydrophobic, often leucine rich, NES^42,45^ that are often found in disordered regions of the cargo proteins. As previously mentioned, NetNES suggested the presence of a conserved NES between V286 and L297 while a Hidden Markov Model predicted that the NES encompassed the residues between L282 and L297 in α-TAT1 C-terminus (Supplementary Fig S2). Interestingly, this region is also predicted to be a site of protein-protein interactions by ANCHOR2 prediction software^40^ (Supplementary Fig. S1b). Truncation of this putative NES, α-TAT1-ΔNES, abrogated its nuclear exclusion (Fig. 3a-e). Alanine substitution of the hydrophobic residues in the NES predicted by the Hidden Markov Model (Supplementary Fig. S2) significantly reduced nuclear exclusion of α-TAT1 (Fig. 3a-d, Supplementary Fig. S6a, b, c), suggesting that these residues contribute to nuclear export of α-TAT1. Taken together, our data suggest that α-TAT1 has a hydrophobic NES in its C-terminus.

### α-TAT1 has a phospho-inhibited NLS in its C-terminus

Our observations thus far with nuclear exclusion of mVenus-α-TAT1 indicate that the putative NLS (P^316^AQRRRT^322^) identified by the PSORT prediction server is either non-functional or is basally inhibited with occasional activation. NLS mediated nuclear import is often phospho-regulated^48^. To identify putative phospho-sites in α-TAT1, we ran its amino-acid sequence through the NetPhos prediction server, which identified over 30 putative phospho-sites in α-TAT1. Nine of these residues have been reported to be phosphorylated in phospho-proteomic studies (Supplementary Fig. S7a, b). Since α-TAT1(1-284) did not display nuclear exclusion (Fig. 3b-e), we reasoned that the phospho-sites which inhibit nuclear localization of α-TAT1 might be located between F285 and R323. NetPhos prediction server identified S294, T303, S315 and T322 as potential phospho-sites in this region (Supplementary Fig. S7a, indicated in red box), wherein only S315 and T322 have been reported to be phosphorylated in phospho-proteomic studies (Supplementary Fig. S7b) and they also flank the putative NLS (Fig. 4a). Importin-α binds with NLS enriched in basic residues through a charge-based interaction^44^. Phosphorylation of amino acids adjacent to such an NLS may inhibit the association of Importin-α binding through a disruption of the charge balance in the NLS region^49^. Alanine substitution of T322, but not of S315, significantly increased nuclear localization of mVenus-α-TAT1 (Fig. 4a-e). Alanine substitution of both S315 and T322, α-TAT1 (ST/A), showed considerably more nuclear localization of mVenus-α-TAT1 than T322 alone (Fig 4a-e). These data suggest that T322 phosphorylation inhibits nuclear localization of α-TAT1, while S315 may play a co-operative role in such inhibition. Substitution of S315 with acidic residues (S315D) appeared to boost nuclear exclusion of α-TAT1, whereas substitution of T322 with acidic residues (T322E) or both (ST/DE) displayed increased diffused pattern, but not increased nuclear accumulation (Supplementary Fig. S8a-c). This may be because these acidic residues, unlike phosphate moieties, do not sufficiently counter the basic residues in the NLS; or that phospho-T322, and to a lesser extent, phospho-S315 phosphorylation may be involved in protein-protein interactions, such as with 14-3-3 adaptor proteins, which inhibit nuclear localization of α-TAT1. These observations suggest that the phosphate moiety in phosphorylated T322 and S315 is critical for inhibition of α-TAT1 nuclear import.

**Figure 4.**
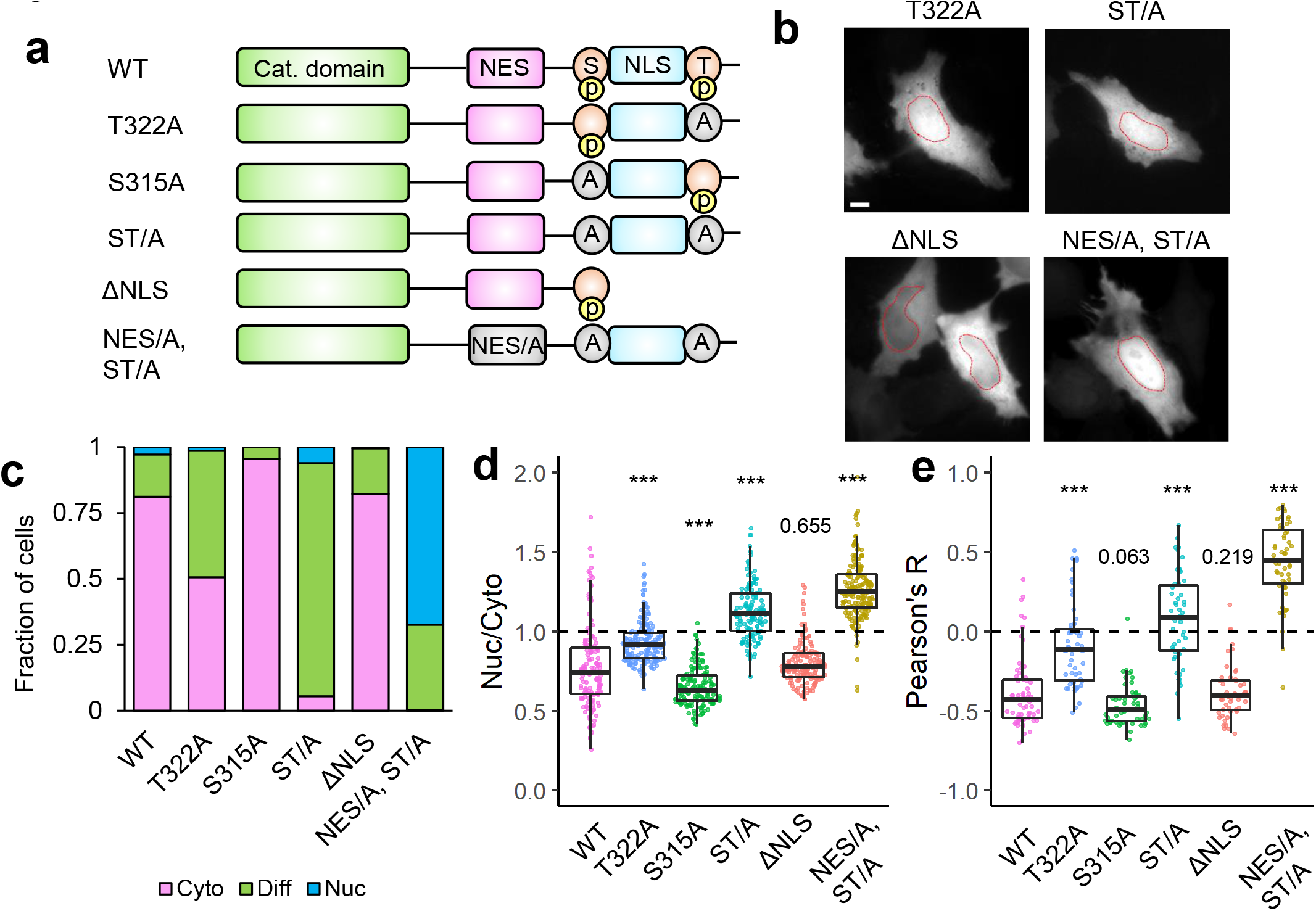
α-TAT1 has a C-terminal phospho-inhibited NLS. a) Cartoon showing α-TAT1 mutant design, b) intracellular distribution of mVenus-α-TAT1 mutants as indicated, nuclei are outlined in red dotted lines, c) categorical analysis (WT: 212, T322A: 246, S315A: 197, ST/A: 190, δNLS: 163, NES/A, ST/A: 186 cells), d) ratiometric analysis (WT: 135, T322A: 170, S315A: 146, ST/A: 137, δNLS: 162, NES/A, ST/A: 195 cells) and e) colocalization analysis(WT: 63, T322A: 54, S315A: 53, ST/A: 51, δNLS: 51, NES/A, ST/A: 55 cells) of intracellular localization of mVenus-α-TAT1 mutants. Scale bar = 10μm, P-value: *** <0.001 or as shown, Student’s *t*-test.

One possible explanation of increased nuclear localization of T322A mutant is that phospho-T322 mediates nuclear export of α-TAT1. Truncation of the putative NLS including T322, α-TAT1-ΔNLS, did not increase nuclear localization (Fig. 4a-e), indicating that T322 did not mediate nuclear export of α-TAT1. Furthermore, alanine substitution of the hydrophobic residues in α-TAT1 NES as well as S315 and T322, α-TAT1(NES/A, ST/A) not only abrogated nuclear exclusion, but considerably increased nuclear accumulation of α-TAT1 (Fig. 5a-d), suggesting additive effects of NES inhibition and NLS activation. α-TAT1(NES/A, ST/DE) mutant also showed diffused pattern but not nuclear accumulation (Supplementary Fig. S8a-c). Taken together, our observations suggest that phospho-T322 inhibits α-TAT1 NLS and that the α-TAT1 NES and NLS act independently of one another.

**Figure 5.**
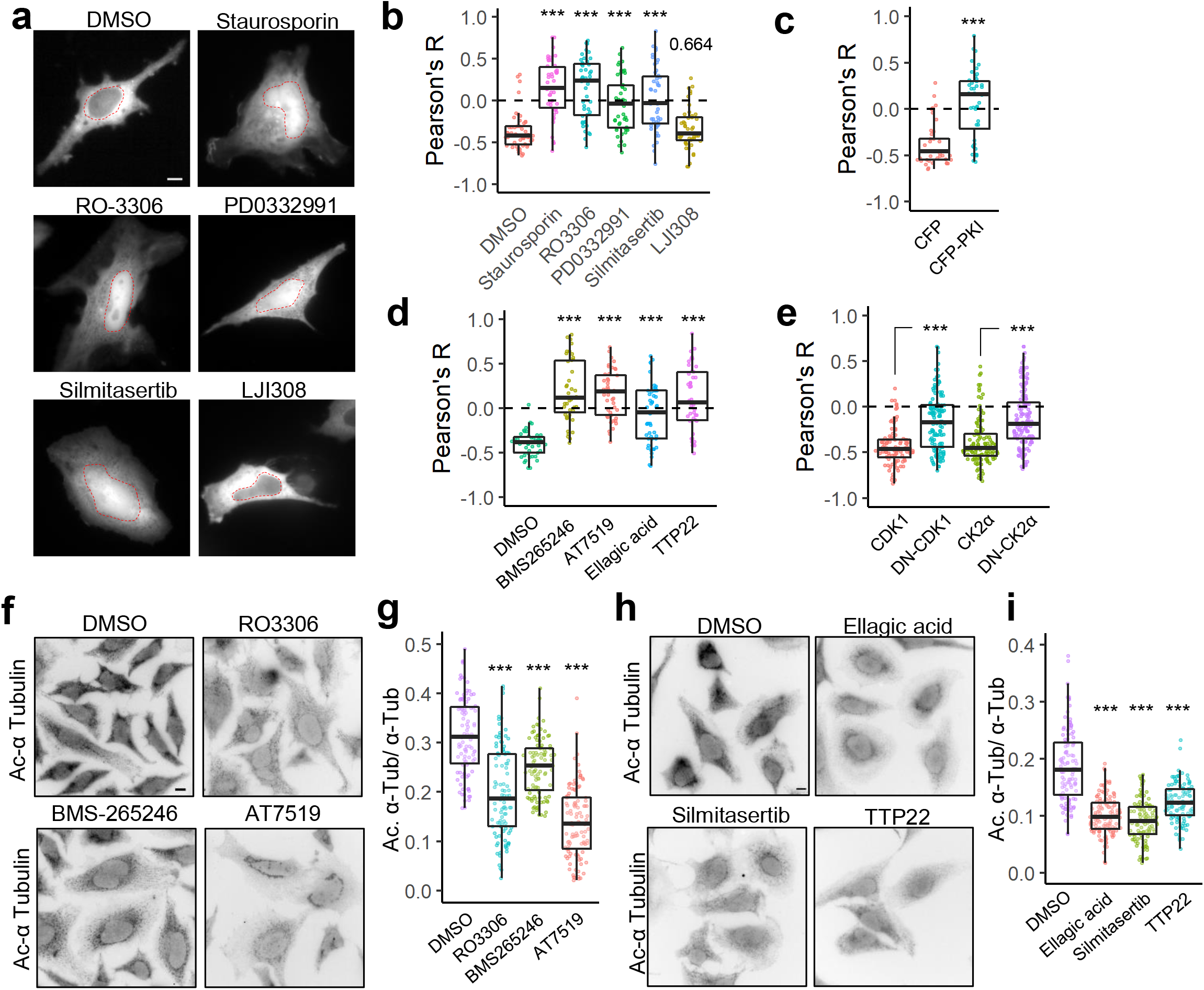
CDK1 and CK2 mediate α-TAT1 localization and MT acetylation. a) Intracellular distribution of mVenus-α-TAT1 in cells treated with DMSO (vehicle) or kinase inhibitors as indicated, nuclei are outlined in red dotted lines, colocalization analyses of intracellular localization of mVenus-α-TAT1 in cells b) treated with DMSO (52 cells), Staurosporine (46 cells), RO-3306 (47 cells), PD0332991 (47 cells), Silmitasertib (47 cells) and LJI308 (46 cells), c) co-expression of CFP (41 cells) or CFP-PKI (44 cells), d) treated with DMSO (44 cells), BMS265246 (43 cells), AT7519 (45 cells), Ellagic acid (52 cells) or TTP22 (42 cells) and e) co-expression of CDK1-mCherry (100 cells), CDK1(D146N)-mCherry (121 cells), mCherry-CK2α (112 cells) and mCherry-CK2α(K68A) (141 cells), f) immunofluorescence images showing acetylated α-tubulin and g) ratio of acetylated to total α-tubulin in HeLa cells treated with vehicle (105 cells), RO-3306 (101 cells), BMS265246 (103 cells) and AT7519 (104 cells), h) immunofluorescence images showing acetylated α-tubulin and i) ratio of acetylated to total α-tubulin in HeLa cells treated with vehicle (114 cells), Ellagic acid (120 cells), Silmitasertib (105 cells) and TTP22 (108 cells). Scale bar = 10μm. P-value: *** <0.001 or as shown, Student’s *t*-test.

### Serine/threonine kinase activities promote cytosolic localization of α-TAT1 and MT acetylation

Since our data suggest that the α-TAT1 NLS is phospho-inhibited, we hypothesized that α-TAT1 localization may be mediated by serine-threonine kinases. Treatment with 100 nM Staurosporin, a pan-kinase inhibitor, significantly increased nuclear localization of α-TAT1 (Fig. 4b, c, Supplementary Fig. S8), suggesting that phosphorylation negatively regulated nuclear localization of α-TAT1. To narrow down the specific kinases that may regulate α-TAT1 localization, we performed a preliminary kinase-inhibitor screening assay where we treated HeLa cells expressing mVenus-α-TAT1 with several serine-threonine kinase inhibitors (summarized in Supplementary Table T1). Based on this screen we identified RO-3306 (inhibitor for CDK1 and CDK2^50,51^), PD0332991 (also called Pablociclib, inhibitor for CDK4 and CDK6^52,53^), Silmitasertib (also called CX-4945, inhibitor for CK2^54^) and mCFP-PKI (peptide-based inhibitor for PKA^55^) as potential inhibitors of nuclear exclusion of mVenus-α-TAT1 (Supplementary Fig. S9a, b, Supplementary Table T1). In addition, LJI308 (inhibitor for Ribosomal S9 kinase) appeared to promote nuclear exclusion of mVenus-α-TAT1 (Supplementary Fig. S9a, b, Supplementary Table T1). The preliminary screening assays were not performed in parallel, and to validate our initial observations, we repeated the kinase inhibition assay with these inhibitors in parallel. Unlike DMSO (vehicle), treatment with Staurosporin, RO-3306, PD0332991 and Silmitasertib significantly increased nuclear localization of mVenus-α-TAT1 (Fig. 5a, b). Colocalization analyses did not show an increase in nuclear exclusion of mVenus-α-TAT1 in cells treated with LJI308. Compared to co-expression with mCFP, mCFP-PKI considerably increased nuclear localization of mVenus-α-TAT1. These data demonstrate that nuclear localization of α-TAT1 is negatively regulated by CDKs, CK2 and PKA. To examine the specificity of kinases, we further treated HeLa cells with exogenous expression of mVenus-α-TAT1 with alternate pharmacological inhibitors BMS-265246 (inhibitor for CDK1 and CDK2), AT7519 (inhibitor for CDK1, CDK2, CDK4, CDK6 and CDK9), Ellagic acid (inhibitor for CK2) and TTP22 (inhibitor for CK2), all of which increased nuclear localization of mVenus-α-TAT1 (Fig. 5d). Additionally, unlike exogenous expression of wild type CDK1 and CK2α, dominant negative Cdk1(D146N) or dominant negative CK2α(K68A) increased nuclear localization of mVenus-α-TAT1 (Fig. 5e, Supplementary Fig. S10).

As exogenous expression of nuclear localized catalytic domain of α-TAT1 had failed to induce MT acetylation (Fig. 1g, h), we reasoned that pharmacological inhibition of CDKs and CK2 may inhibit its function and reduce MT acetylation. HeLa cells treated with CDK1 inhibitors (RO-3306, BMS-265246 or AT7519) or CK2 inhibitors (Ellagic acid, Silmitasertib or TTP22) showed significantly reduced MT acetylation levels unlike those treated with DMSO (vehicle) (Fig. 5f-i). Taken together, these data demonstrate that inhibition of CDK1 (and potentially other CDKs) and CK2 increase nuclear localization of mVenus-α-TAT1 and inhibit MT acetylation.

### α-TAT1 interacts with 14-3-3 proteins

14-3-3 protein binding has been reported to negatively regulate nuclear import by inhibiting binding of importins to NLS^56–58^. 14-3-3 binding to proteins is mediated by phosphorylated serine and threonine residues^59^. Part of the putative NLS sequence “PAQRRRTR” bears similarity to 14-3-3 binding motif RXX(pS/pT)XP^59^, and 14-3-3-Pred^60^, a 14-3-3 interaction prediction server identified T322 as a potential 14-3-3 binding site. In a previous mass spectrometry analysis of α-TAT1 in HEK cells, we had identified 14-3-3-β and 14-3-3-ζ as potential interactors^61^. To examine whether α-TAT1 bound to 14-3-3 proteins, we co-expressed GFP-α-TAT1 with HA-tagged 14-3-3-β or 14-3-3-ζ in HEK-293T cells and performed a co-immunoprecipitation assay. GFP-α-TAT1, but not GFP alone, co-precipitated with HA-14-3-3-β and HA-14-3-3-ζ (Fig. 6a). Exogenous expression of a 14-3-3 inhibitor peptide R18^62^ (PHCVPRDLSWLDLEANMCLP) significantly increased nuclear localization of mVenus-α-TAT1 in HeLa cells (Fig. 6b, c). To further identify which 14-3-3 isoforms interacted with α-TAT1, we performed a chemical-induced dimerization-based protein-protein interaction assay in HeLa cells (see Methods, Supplementary Fig. S11a). Based on this assay, we found that mVenus-FKBP-α-TAT1 interacted with all seven isoforms of 14-3-3 proteins (Fig. 6d, f, h, Supplementary Fig. S11b). To further examine whether any 14-3-3 isoform specifically interacted with T322 and S315 in α-TAT1, we performed the protein-protein interaction assay with mVenus-FKBP-α-TAT1(T322A) and mVenus-FKBP-α-TAT1 (ST/A), both of which showed significantly decreased interaction with 14-3-3 isoforms β, γ, ε and ζ, but not η, θ or σ (Fig. 6e, g, h, Supplementary Fig. S11b). These data suggest that α-TAT1 interacts with specific 14-3-3 isoforms through T322, and that 14-3-3 proteins mediate nuclear exclusion of α-TAT1. Based on these observations, we propose a model of spatial regulation of α-TAT1 wherein a balance of phospho-regulated nuclear transport and 14-3-3 association mediates α-TAT1 localization and MT acetylation (Fig. 6i).

**Figure 6.**
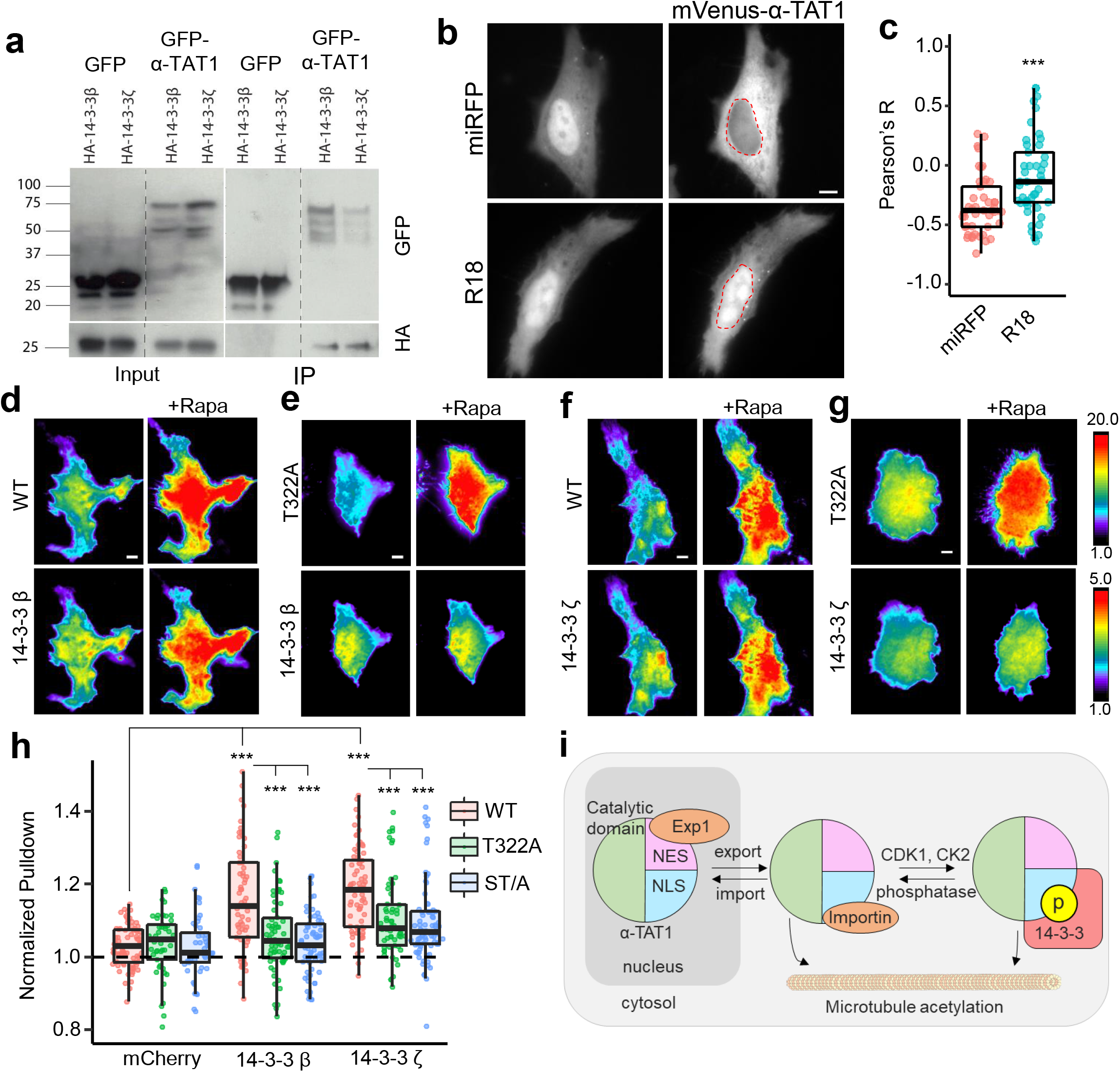
14-3-3 proteins interact with α-TAT1 through T322. a) Co-immunoprecipitation of GFP-α-TAT1 with HA-14-3-3β and HA-14-3-3ζ proteins, b) intracellular distribution of mVenus-α-TAT1 in cells co-expressing miRFP703 or miRFP703-14-3-3i, nuclei are outlined in red dotted lines, c) colocalization analysis of mVenus-α-TAT1 in cells co-expressing miRFP703 (45 cells) or miRFP703-14-3-3i (45 cells), d) images showing changes in TIRF intensity in CID based protein-protein interaction assay for mCherry-14-3-3β with mVenus-FKBP-α-TAT1 and e) mVenus-FKBP-α-TAT1 (T322A), f) mCherry-14-3-3ζ with mVenus-FKBP-α-TAT1 or with g) mVenus-FKBP-α-TAT1 (T322A), h) Normalized co-recruitment levels by indicated baits of mCherry (α-TAT1: 79, T322A: 45, ST/A: 49), mCherry-14-3-3ζ (α-TAT1: 70, T322A: 58, ST/A: 79), (complete panel: Supplementary Fig. S11b), i) proposed model of spatial regulation of α-TAT1 function. Scale bar = 10μm. P-value: *** <0.001 or as shown, Student’s *t*-test.

### Phospho-regulated intracellular distribution of α-TAT1 mediates MT acetylation

To further validate our proposed model, we utilized mouse embryonic fibroblasts (MEFs) isolated from α-TAT1 knock-out (KO) mice^63^. Compared to MEFs isolated from wild-type (WT) mice, KO MEFs showed significantly lower levels of acetylated MTs (Fig. 7a, Supplementary Fig. S12a). Additionally, pharmacological inhibition of histone deacetylase-6 (HDAC-6), a known deacetylase for α-Tubulin, increased MT acetylation in WT MEFs but not KO MEFs (Fig. 7a, Supplementary Fig. S12), further demonstrating that α-TAT1 is the primary MT acetyltransferase in mice. To examine our proposed model, we used lentiviral transduction of the KO cells to stably express mVenus, mVenus-α-TAT1, mVenus-α-TAT1(ST/A) and mVenus-α-TAT1 (NES/A, ST/A), which were further selected using flow cytometry to isolate cell populations exhibiting similar levels of mVenus fluorescence to ensure comparable expression levels of the introduced protein. Consistent with our previous observations, mVenus-α-TAT1 was predominantly excluded from the nuclei, whereas mVenus, mVenus-α-TAT1 (ST/A) showed diffused pattern indicating a loss of nuclear exclusion and mVenus-α-TAT1 (NES/A, ST/A) showed nuclear enrichment (Supplementary Fig. S13). Rescue with mVenus-α-TAT1, but not mVenus, restored MT acetylation in KO MEFs, but at levels significantly below those of WT MEFs treated with tubacin. Moreover, KO MEFs expressing mVenus-α-TAT1 (ST/A) or mVenus-α-TAT1 (NES/A, ST/A) showed reduced levels of MT acetylation (Fig. 7a, Supplementary Fig. S12), validating our proposed model.

**Figure 7.**
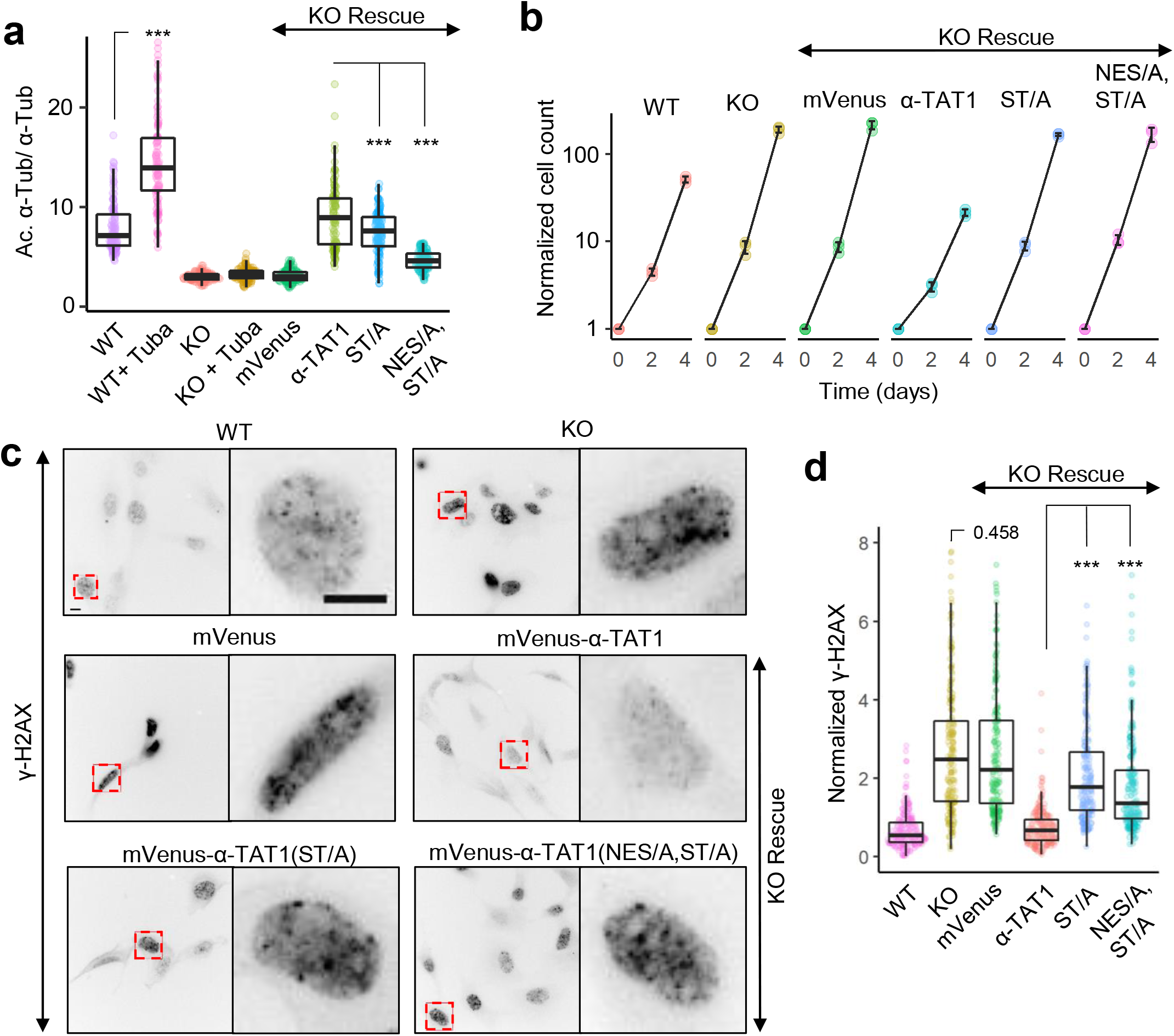
Phosphoregulated spatial distribution of α-TAT1 mediates MT acetylation, cell proliferation and cellular DNA damage response. a) Ratio of acetylated to total α-tubulin in MEFs (WT: 110, KO: 120, WT + tubacin: 110, KO + tubacin: 116, mVenus: 110, mVenus-α-TAT1: 100, mVenus-α-TAT1(ST/A): 132 and mVenus-α-TAT1(NES/A,ST/A): 112 cells), b) normalized changes in cell number over 48 and 96 hours in WT, KO and KO-rescue cells as indicated (n = 3 for each sample, individual points and standard deviation are shown), c) γ-H2AX intensity levels (WT: 214, KO: 214, mVenus: 187, mVenus-α-TAT1: 226, mVenus-α-TAT1 (ST/A): 217 and mVenus-α-TAT1 (NES/A,ST/A): 223 cells) and d) immunofluorescence images showing γ-H2AX levels in in WT and KO and KO-rescue MEFs as indicated, inset red boxes are shown in the adjacent panel. Scale bar = 10μm. P-value: *** <0.001 or as shown, Student’s *t*-test.

### Cytoplasmic localization of α-TAT1 mediates cell proliferation and DNA damage response

To understand whether spatial regulation of α-TAT1 has any pertinence in cell behaviour, we examined cell proliferation and DNA damage response in cells. In line with previous reports^63^, KO MEFs showed increased proliferation compared to WT MEFs (Fig. 7b). KO MEFs expressing mVenus-α-TAT1, but not mVenus, mVenus-α-TAT1(ST/A) or mVenus-α-TAT1 (NES/A, ST/A) showed a significant reduction in proliferation rates (Fig. 7b). CDK1 activity is maximal between G1 and M phase in the cell cycle^64^, towards the end of which, (G2/M transition) there is breakdown of the nuclear envelope, which would render spatial regulation of α-TAT1 redundant. We reasoned that the cell cycle defects due to loss of spatial regulation of α-TAT1 may be due to events in G1 or S phase, where CDK1 is active and nuclear envelope is intact. Both CDK1 and CK2 have been implicated in DNA damage response in cells^64–66^. α-TAT1 depleted HeLa cells demonstrated defects in cell cycle arrest after DNA damage^67^. Immunostaining for γ-H2AX, a marker for DNA damage, showed significantly higher intensity in KO MEFs compared to WT (Fig. 7c, d), suggesting that α-TAT1 or MT acetylation was critical for cellular response to DNA damage. KO-MEFs expressing mVenus-α-TAT1, but not mVenus, mVenus-α-TAT1 (ST/A) or mVenus-α-TAT1 (NES/A, ST/A) restored DNA damage response to levels comparable to WT MEFs (Fig. 7c, d). These observations confirm a role of phospho-regulated nuclear transport of α-TAT1 in its function and suggest a role of MT acetylation, or that of α-TAT1 C-terminus in DNA damage response and cell cycle progression.

## Discussion

One of the bottlenecks in elucidating the role of microtubule acetylation in biological phenomena is the knowledge gap of how upstream molecular signaling pathways control α-TAT1 function to modulate MT acetylation. Auto-acetylation of lysine residues is proposed to promote α-TAT1 catalytic activity^38^. Similarly, TAK1 dependent phosphorylation of α-TAT1 Serine-237 has been reported to stimulate its catalytic property^68^. In neurons, p27^kip1^ directly binds to α-TAT1 and stabilizes it against proteasomal degradation^24^, thus enhancing MT acetylation. Our study demonstrates that intracellular α-TAT1 localization is a dynamic process that is orchestrated on a delicate balance of nuclear export and import, and that is directly reflected in MT acetylation levels (Fig. 6i). Based on decrease in MT acetylation in the presence of kinase inhibitors or KO-rescue with α-TAT1 (NES/A, ST/A), we posit that the effects of spatial regulation are at least comparable to previously reported regulatory mechanisms^24,38,68^, suggesting a significant role in tuning MT acetylation levels. To our knowledge, this is the first study to identify the molecular mechanisms that spatially regulate α-TAT1 function. In addition, we demonstrated a hitherto unknown role of the inherently disordered α-TAT1 C-terminus and identified its interactions with specific 14-3-3 proteins and serine-threonine kinases, namely CDK1 and CK2. Our observation that spatial sequestration of α-TAT1 from cytosolic MTs modulates acetylation dynamics suggests a role of the nucleus as a reservoir or sequestration chamber to control protein access of substrates.

We have demonstrated active nuclear export of α-TAT1 by Exp1 through an NES rich in hydrophobic residues, which was critical for efficient microtubule acetylation. In addition, we have identified an NLS consistent with non-canonical class IV NLS^44^. Interestingly, position 7 of this NLS, which should not be an acidic residue, is occupied by Threonine-322. Since phosphorylation of threonine can significantly increase its net negative charges, it is ideally situated to act as an ON/OFF switch for the NLS. Although we have identified Threonine-322 to be the critical phospho-residue that inhibits nuclear import, Serine-315 appears to provide additional inhibition. The increased nuclear localization of ST/A mutant over T322A mutant raises the possibility that S315 and T322 may aggregate signals from different signaling pathways to fine-tune α-TAT1 localization.

Our data demonstrate that cytosolic localization of α-TAT1 is facilitated by kinase action, possibly on Threonine-322 and Serine-315. Specifically, our study shows a role of CDK1 and CK2 in coordinating spatial distribution of α-TAT1 and downstream MT acetylation to mediate cell proliferation and DNA damage response. Such phospho-regulation of α-TAT1 provides a possible mechanism for the changes in α-TAT1 localization and microtubule acetylation observed at different stages of the cell cycle^69^. While MT acetylation plays a critical role in DNA damage response, whether Threonine-322 mediates DNA damage response through modulating MT acetylation levels or by additional means remains to be examined. We demonstrated Threonine-322 to be a putative binding site for 14-3-3 β, γ, ε and ζ proteins, which are also involved in cellular DNA damage response and cell cycle progression^70–72^. 14-3-3s interact with phospho-serines or phospho-threonines in intrinsically disordered regions and may mediate nuclear transport of proteins by masking NES or NLS^72^. Furthermore, 14-3-3 proteins may significantly alter the structure of their binding partners to align along their rigid α-helical backbone, to expose or hide critical binding sites^73^.

While there are pharmacological agents that promote increased MT acetylation through inhibition of deacetylases, currently there are no available pharmacological inhibitors of α-TAT1 itself, although there appears to be a considerable interest in the development of such inhibitors^74^. Identifying a small chemical targeted to the C-terminus signal motif to alter the subcellular localization, instead of the catalytic activity, may open up a novel approach for inhibiting MT acetylation. It is possible that nuclear import of α-TAT1 facilitates interactions with presently unidentified substrates located in the nucleus, or that the C-terminus of α-TAT1 facilitates other protein-protein interactions. In a similar vein, lysine residues in cyclins, CDK1, CK2, 14-3-3 proteins and Exp1 are acetylated^75–77^, and it is tempting to speculate that these acetylation events may be directly or indirectly mediated by α-TAT1.

It is worthwhile to consider that a significant number of post-translational modifications of α-TAT1 appear on its intrinsically disordered C-terminus (Supplementary Fig. 7b). Disordered regions may act as a signaling hub by interacting with multiple proteins, thus facilitating complex formation and acting as integrators of signaling pathways^78^. We demonstrated the presence of an NES, NLS, phosphorylation sites and putative 14-3-3 binding sites within the α-TAT1 C-terminus. This signal motif is well conserved across mammalian species as well as in all the human isoforms, suggesting a critical role of the α-TAT1 C-terminus in its function. The inability of the α-TAT1(ST/A) to rescue the defects in the cellular DNA damage response and increased cell proliferation state suggests a potential role of the α-TAT1 C terminus in cancer. Indeed, numerous cancer-associated mutations curated in COSMIC^79^ and TCGA Research Network databases in the ATAT1 gene are located in the intrinsically disordered C-terminal region. More specifically, there are a considerable number of deletions, frame shifts and missense mutations encompassing the NES and the NLS regions, which may be expected to affect the spatial distribution of α-TAT1. Whether these mutations underlie the pathogenesis in these cancers remain to be examined. Considering the role of microtubule acetylation in a wide array of cellular activities, it may be conjectured that loss of spatial regulation of α-TAT1 may be present in other diseases as well.

Kinase-mediated regulation of nuclear export and nuclear import has previously been reported in transcription regulators^80–83^. In particular, regulation of Cdc25 localization by Checkpoint kinase1 (Chk1) mediated phosphorylation of and subsequent recruitment of 14-3-3-β to an NLS-proximal phospho-site is virtually the same as our proposed model (Fig. 6i) of α-TAT1 localization^83–85^, suggesting that such kinase-mediated balancing of nuclear export and import is a general strategy for protein localization. This is particularly intriguing in the context of the apparent role of α-TAT1 or MT acetylation in DNA damage response since Chk1 association with 14-3-3 is triggered by DNA damage^71,86^. Localization of proteins is often regulated^87–93^ and their aberrant regulation have been linked to diseases^94–97^. For example, class IIa histone deacetylases (HDACs) may be accumulated in the cytosol by 14-3-3 proteins, which inhibits acetylation of transcription factors^93^. Cytosolic retention of p42/44 MAP kinases by PEA-15 inhibits its effects on transcription and proliferation^89^. Unlike these proteins, the nuclear localization of α-TAT1 limits the access of the enzyme to its substrates, namely MTs, which exclusively reside in the cytosol.

In conclusion, we propose a model for spatial sequestration of α-TAT1 as the major regulatory mechanism of microtubule acetylation (Fig. 6i). The model consists of three key characteristics: presence of an NES that facilitates Exp1 mediated nuclear export, presence of an NLS to mediate nuclear import and finally, modulation of this nuclear import by kinases. Further examination of the role of specific kinases on α-TAT1 localization may advance our understanding of its function in both cellular processes and pathologies, helping identify new therapeutic targets in the future.

## Materials and Methods

### Cell culture and transfection

HeLa and HEK-293T cells were cultured in DMEM basal media and passaged every third day of culture. For optimal growth, the media were supplemented with 10% (v/v) fetal bovine serum, L-Glutamine and Penicillin/Streptomycin. WT and α-TAT1 KO MEFs were a generous gift from Dr. Maxence Nachury and were cultured in DMEM basal media supplemented with 10% (v/v) fetal bovine serum, L-Glutamine, Penicillin/Streptomycin, Non-essential amino acids and 0.05 mM β-mercaptoethanol. KO MEFs rescues with mVenus, mVenus-α-TAT1, mVenus-α-TAT1 (ST/A) or mVenus-α-TAT1 (NES/A, ST/A) were selected using the Sony SH800 cell sorter using manufacturer’s instructions to select cell populations with the same mVenus fluorescence thresholds to ensure similar expression of the proteins of interest. The cells were maintained under standard cell culture conditions (37 °C and 5% CO_2_) and were checked for mycoplasma contamination prior to use in experiments. In addition, the KO-rescue cells were cultured in medium containing 1 μg/ml of puromycin for selection; effective puromycin dosage was ascertained by testing on KO cells. FuGENE 6 reagent (Promega, Madison, WI) was used for transient transfection of HeLa cells according to the manufacturer’s instructions. For immunoprecipitation assays and generation of lentiviral particles, HEK cells were transfected using polyethyleneimine (PEI) method.

### DNA plasmids

H2B-m Cherry construct was a generous gift from Dr. Sergi Regot. α-TAT1 construct was a generous gift from Dr. Antonina Roll-Mecak. The α-TAT1 construct was subcloned into the pTriEx-4 vector (Novagen) using PCR and restriction digestion with mVenus at the N terminus and α-TAT1 at the C terminus. H2B-mCherry and CFP-PKI constructs were respectively subcloned into mCherry-C1 and mCer3-C1 vectors (Clontech). GFP-αTAT1 construct was a gift from Dr. Philippe Chavrier and Dr. Guillaume Montagnac. HA-14-3-3 plasmids were a generous gift from Dr. Michael Yaffe. pRS3 (CK2alpha, CK2beta) was a gift from David Litchfield (Addgene plasmid # 27092). pUHD Cdk1 WT HA was a gift from Greg Enders (Addgene plasmid # 27652). CK2-α and CDK1 were sub-cloned into pTriEx-4 vector. As indicated in the results and figure legends, tags of compatible fluorescent proteins including Cerulean, mVenus and mCherry were appended to facilitate detection. Unless specified otherwise, the termini of tagging were positioned as in the orders they were written. Lentiviral plasmids were generated based on a modified Puro-Cre vector (Addgene plasmid # 17408, mCMV promoter and no Cre encoding region). Truncations of α-TAT1 were generated by PCR. Point mutations of α-TAT1 were generated using overlapping PCR. The open reading frames of all DNA plasmids were verified by Sanger sequencing.

### Sequence alignment

Protein sequence alignment was performed using Clustal-W^98^ (https://www.ebi.ac.uk/Tools/msa/clustalo/).

### Nuclear transport and kinase inhibitors

LMB was purchased from LC Laboratories (catalog # L6100). SB203580 (Sigma Aldrich, catalog # S8307), Doramapimod (BIRB 796, Selleck Chemicals, catalog # S1574), CHIR99021 (Sigma Aldrich, catalog # SML1046), Sostrastaurin (Selleck Chemicals, catalog # S2791), RO-3306 (Selleck Chemicals, catalog # S7747), Ipatasertib (RG7440, Selleck Chemicals, catalog # S2808), Capivasertib (AZD5363, Selleck Chemicals, catalog # S8019), Silmitasertib (CX 4945, Selleck Chemicals, catalog # S2248), KU-55933 (Sigma Aldrich, catalog # SML1109) were generous gifts from Dr. Sergi Regot. SB239063 (Sigma Aldrich, catalog # S0569) was a generous gift from Dr. Jun Liu. The rest of the kinase inhibitors were purchased as indicated: Staurosporine (Sigma Aldrich, catalog # 569397), LJI308 (Sigma Aldrich, catalog # SML1788), Y-27632 (LC Laboratories, catalog # Y-5301), Gö 6976 (Sigma Aldrich, catalog # 365250), Gö 6983 (Sigma Aldrich, catalog # G1918), H-89 (Sigma Aldrich, catalog # B1427), D4476 (BioVision, catalog # 1770), KN-62 (Selleck Chemicals, catalog # S7422), KU-57788 (MedChemExpress, catalog # HY-11006), Purvalanol B (AdipoGen Life Sciences, catalog # SYN-1070), KT-5823 (Cayman Chemicals, catalog # 10010965), PD0332991 (Sigma Aldrich, catalog # PZ0199), BMS-265246 (Selleck Chemicals, catalog # S2014), AT7519 (Selleck Chemicals, catalog # S1524), TTP22 (Selleck Chemicals, catalog # S6536) and Ellagic acid (Selleck Chemicals, catalog # S5516).

### Immunofluorescence assays

For immunostaining of acetylated and total α-Tubulin, cells were fixed using ice-cold methanol for 10 minutes, washed thrice with cold PBS, blocked with 1% BSA in PBS for one hour and then incubated overnight at 4°C with monoclonal antibodies against tubulin (Millipore, catalog # MAB1864) and acetylated α-Tubulin (Sigma Aldrich, catalog # T7451). Next day, the samples were washed thrice with cold PBS and incubated with secondary antibodies (Invitrogen) for one hour at room temperature, after which they were washed thrice with PBS and images were captured by microscopy. For HeLa cells transiently transfected with mVenus-α-TAT1, mVenus-α-TAT1 catalytic domain and NLS-mVenus-α-TAT1 catalytic domain, fixation and immunostaining were performed 24 hours post-transfection. For LMB treatment, HeLa cells were treated with 100 nM LMB or equal volume of vehicle (EtOH) incubated for 4 hours, followed by methanol fixation and immunostaining. For kinase inhibitors, HeLa cells were treated with RO-3306 (10 μM), BMS-265246 (100 nM), AT7519 (1 μM), Ellagic acid (1 μM), TTP22 (1 μM) or Silmitasertib (10 μM), or vehicle (DMSO) for 4 hours followed by fixation and immunostaining. For Tubacin treatment, MEF cells were treated with 2 μM Tubacin or vehicle (DMSO), incubated for 4 hours. KO-rescue cells were treated with DMSO for 4 hours.

For γ-H2AX staining, MEF cells were fixed using 4% paraformaldehyde at room temperature for 15 minutes, washed twice with PBS, permeabilized with 0.1% TritonX-100 in PBS for 5 minutes, washed thrice with PBS, blocked with 1% BSA in PBS for 1 hours at room temperature and then incubated overnight at 4°C with monoclonal antibody against phospho-Histone H2A.X (Ser139) Antibody, clone JBW301 (Sigma-Aldrich, catalog # 05-636-25UG). Next day, the samples were washed thrice with cold PBS and incubated with secondary antibody (Invitrogen) and DAPI for one hour at room temperature, after which they were washed thrice with PBS and images were captured by microscopy.

### Immunoprecipitation assays

HEK293T cells were transiently transfected with pEGFP-c1 (GFP-Ctl) or GFP-αTAT1 with HA-14-3-3β or HA-14-3-3ζ using the calcium phosphate method. Cell lysates were prepared by scraping cells using 1X lysis buffer (10X recipe-50 mM Tris pH 7.5, Triton 20%, NP40 10%, 2 M NaCl, mixed with cOmplete protease inhibitor tablet - Roche, Product number 11873580001). Cell lysates were rotated on a wheel at 4°C for 15 min and centrifuged for 10 min at 13,000 rpm 4°C to pellet the cell debris. A small volume of the supernatant was used as the soluble input. Soluble detergent extracts were incubated with GFP nanobody (NanoTag, N0310) for 1 h at 4°C. Samples were then centrifuged and washed thrice with wash buffer (250 mM NaCl, 0.1% Triton X-100 in PBS). The resin and the soluble input were then mixed with Laemmli buffer (composed of 60 mM Tris-HCl pH 6.8, 10% glycerol, 2% SDS and 50 mM DTT with the addition of protease and phosphatase inhibitors). Samples were boiled 5 min at 95°C before loading in polyacrylamide gels. Gels were transferred for western blot and membranes were blocked with TBST (0.1% Tween) and 5% milk and incubated 1 h with the primary antibody and 1 h with HRP-conjugated secondary antibody. Bands were revealed with ECL chemiluminescent substrate (Biorad). Two different western blots were used to visualize GFP-Input and HA-14-3-3 proteins due to similar molecular weights. Antibodies used: GFP-HRP (NB600-313, Novus Biologicals), anti-HA (rat; Merck; 11867423001), anti-exportin-1 (mouse; BD Transduction Laboratories™; 611832). Secondary HRP antibodies were all purchased from Jackson ImmunoResearch.

### Chemically-inducible co-recruitment assay

mVenus-FKBP was fused to α-TAT1, while FRB is tethered to the inner leaflet of plasma membrane using the CAAX-region of K-Ras. Upon rapamycin addition, FKBP binds to FRB which brings the bait (mVenus-FKBP-α-TAT1) and the prey capable of binding (m Cherry-14-3-3) to the plasma membrane. Recruitment of the bait and the prey to the plasma membrane were detected by TIRF microscopy as an increased fluorescence signal (Supplementary Fig. S11a). For quantification, after background subtraction, co-recruitment levels of prey were measured by increase in mCherry (prey) signal normalized to the intensity before rapamycin addition. Only cells showing at least 30% increase in mVenus (bait) intensity after Rapamycin addition were considered.

### Microscopy and image analyses

All imaging was performed with an Eclipse Ti microscope (Nikon) with a 100X objective (1.0X zoom and 4X4 binning) and Zyla 4.2 sCMOS camera (Andor), driven by NIS Elements software (Nikon). Time-lapse imaging was performed at 15 min intervals for 10-15 hours. All live cell imaging was conducted at 37°C, 5% CO_2_ and 90% humidity with a stage top incubation system (Tokai Hit). Vitamin and phenol red-free media (US Biological) supplemented with 2% fetal bovine serum were used in imaging to reduce background and photobleaching. Inhibitors and vehicles were present in the imaging media during imaging. All image processing and analyses were performed using Metamorph (Molecular Devices, Sunnyvale, CA, USA) and FIJI software (NIH, Bethesda, MD, USA).

For kinase inhibitor assays, cells showing high degree of blebbing were excluded from analysis to minimize artifacts from possible induction of apoptosis. For categorical analysis of mVenus-α-TAT1 localization, images were visually inspected and classified as displaying either cytosolic, diffused, or nuclear enriched localization of mVenus fluorescence signal. For ratiometric analysis, the ratio of the fluorescence intensity from region of interest (≈ 10 μm diameter) inside the nucleus (as identified by H2B-mCherry or Hoechst staining) to that in a perinuclear area was used to minimize any volumetric artifacts (Supplementary Fig. S4a). For all ratiometric analyses, background subtraction based on a cell free area on each image was performed prior to calculation of the ratio. For colocalization analysis, Coloc2 function in FIJI was used to calculate the Pearson’s R coefficient value between the mVenus-α-TAT1 image and the nuclear marker (H2B-Cherry or Hoechst). Pearson’s R value above threshold is reported. Only cells showing at least a signal-to-noise ratio of at least 2 in mVenus, mCherry or DAPI (Hoechst) channel were used (~ 12 bit to 16 bit range). Images containing any saturated pixels (65535 value) within the cell area were excluded.

For immunofluorescence assays with exogenous expression of mVenus-α-TAT1 plasmids, transfected cells were identified by the presence of mVenus fluorescence signal. The ratio of acetylated α-Tubulin over α-Tubulin (Ac. α-Tub/α-Tub) for transfected cells was normalized against that for non-transfected cells averaged over 20 untransfected cells from the same dish. For LMB and vehicle treatment, Ac. α-Tub/α-Tub ratios are shown. For LMB or kinase inhibitor, tubacin treatment or KO-rescue cells, ratio of acetylated α-Tubulin over α-Tubulin (Ac. α-Tub/α-Tub) for individual cells are shown. For γ-H2AX staining, nuclear region was identified using DAPI staining and was used to compute average intensity of nuclear γ-H2AX stain.

### Statistical analyses

Microsoft Excel (Microsoft, Redmond, WA, USA) and R (R Foundation for Statistical Computing, Vienna, Austria) were used for statistical analyses. The exact number of samples for each data set is specified in the respective figure legends. Data was pooled from at least three independent experiments performed in parallel. For the kinase inhibitor screening assay, data were pooled from at least three independent experiments for each inhibitor, but these experiments were not performed in parallel. To validate the preliminary observations from the screening assay, experiments were repeated with the inhibitors of interest with appropriate vehicle controls and performed in parallel. Sample sizes were chosen based on the commonly used range in the field without performing any statistical power analysis. Normal probability plot (Supplementary Fig. S14) was utilized to confirm normal distribution of the Nuc/Cyto ratio or Pearson’s R. *P*-values were obtained from two-tailed Student’s *t*-test assuming equal variance. *P*-values > 0.001 are shown in the figures. Exact *P*-values for kinase inhibitor screening assay are available in Supplementary Table T1.

## Data availability

All data and constructs used in this project are available upon reasonable request.

## Acknowledgements

We thank Allen Kim for discussions that led to initiation of this project, Amy F. Peterson for help with the kinase inhibitor assays and Dipika Gupta for discussions on the DNA damage assays. We thank Yuta Nihongaki and Helen D. Wu for constructive discussions, as well as Robert DeRose for proofreading. This project was supported by American Heart Association fellowship 20POST35220046 (ADR), discretionary funds (TI), the La Ligue contre le cancer (S-CR17017) and Centre National de la Recherche Scientifique and Institut Pasteur. SS is funded by the ITN PolarNet Marie Curie grant and Fondation pour la Recherche Médicale and is enrolled at the Ecole Doctorale Frontières du Vivant (FdV) _ Programme Bettencourt.

## Contributions

ADR initiated the project and designed and performed most of the experiments and data analyses. GSP and EGG performed experiments and data analyses under the guidance of ADR and TI. SS performed the immunoprecipitation assays under the guidance of SEM. ADR and TI wrote the final version of the manuscript based on contributions from all the authors.

