## Supplementary Information for "A phospho-regulated signal motif determines subcellular localization of α-TAT1 for dynamic microtubule acetylation"

MEFPFDVDALFPERITVLDQHLRPPARRPGTTTPARVDLQQQIMTIIDELGKASAKAQNLSAPITSA
SRMQSNRHVVYILKDSSARPAGKGAIIGFIKVGYKKLFVLDDREAHNEVEPLCILDFYIHESVQRHG
HGRELFQYMLQKERVEPHQLAIDRPSQKLLKFLNKHYNLETTVPQVNNFVIFEGFFAHQHRPPAPS
LRATRHSRAAAVDPTPAAPARKLPPKRAEGDIKPYSSSDREFLKVAVEPPWPLNRAPRRATPPAH
PPPRSSSLGNSPERGP**LRPFVPEQELLRSLRL**CPPHPTARLLLAADPGGS$^{315}$**PAQRRRT**$^{322}$R

b)

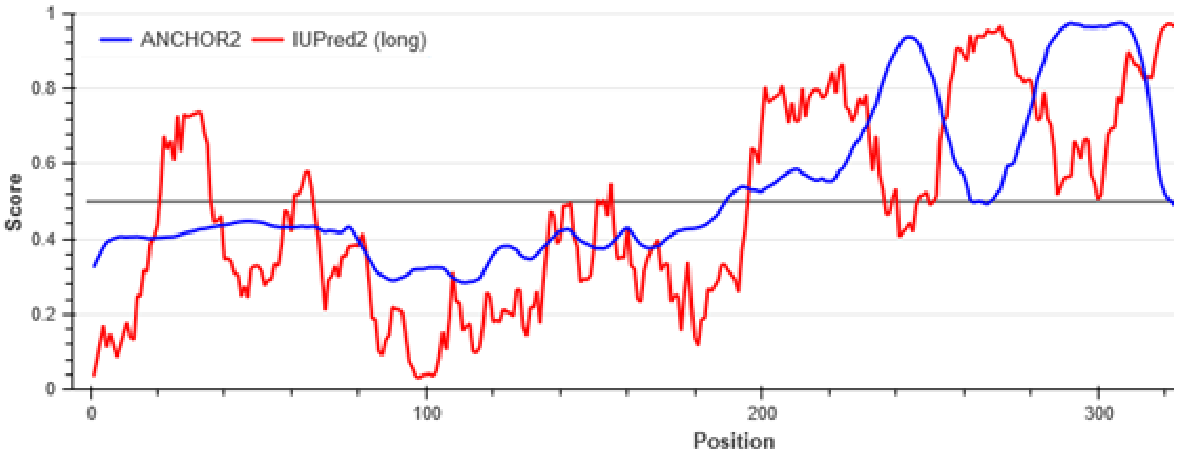


c)

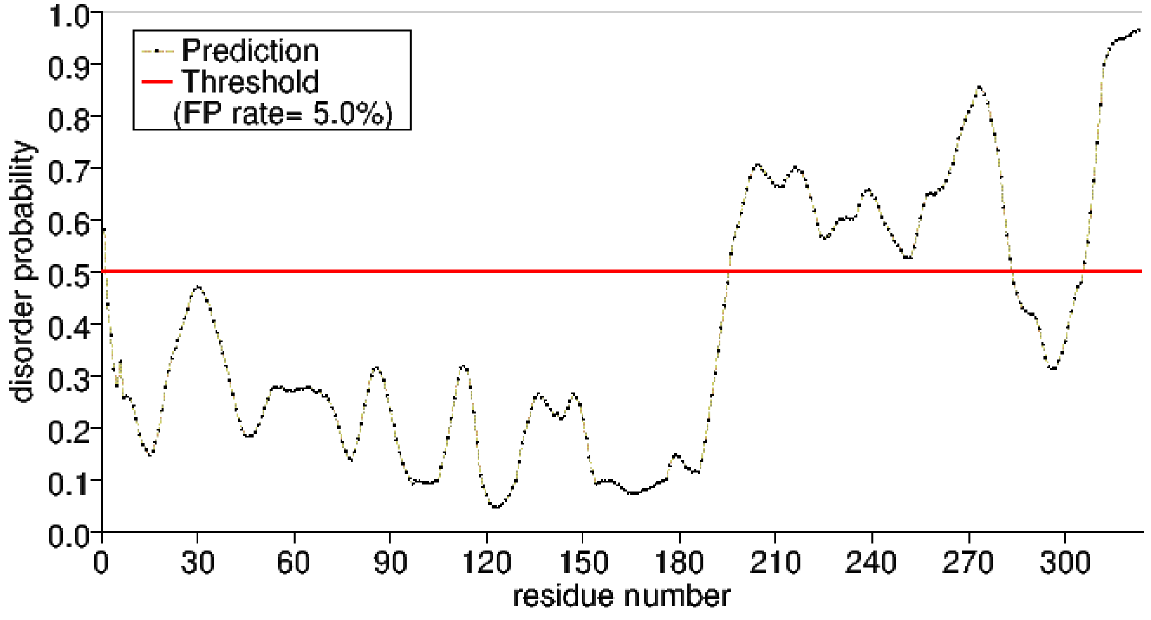


Supplementary Figure S1. αTAT1 C-terminus is disordered. a) amino acid sequence of αTAT1 isoform 7 used as the query, the putative NES and NLS are shown in bold, b) IUPred2 (and ANCHOR2) and c) PrDOS results suggesting that αTAT1 C-terminus is disordered. The threshold to be considered disordered is 0.5 for both.

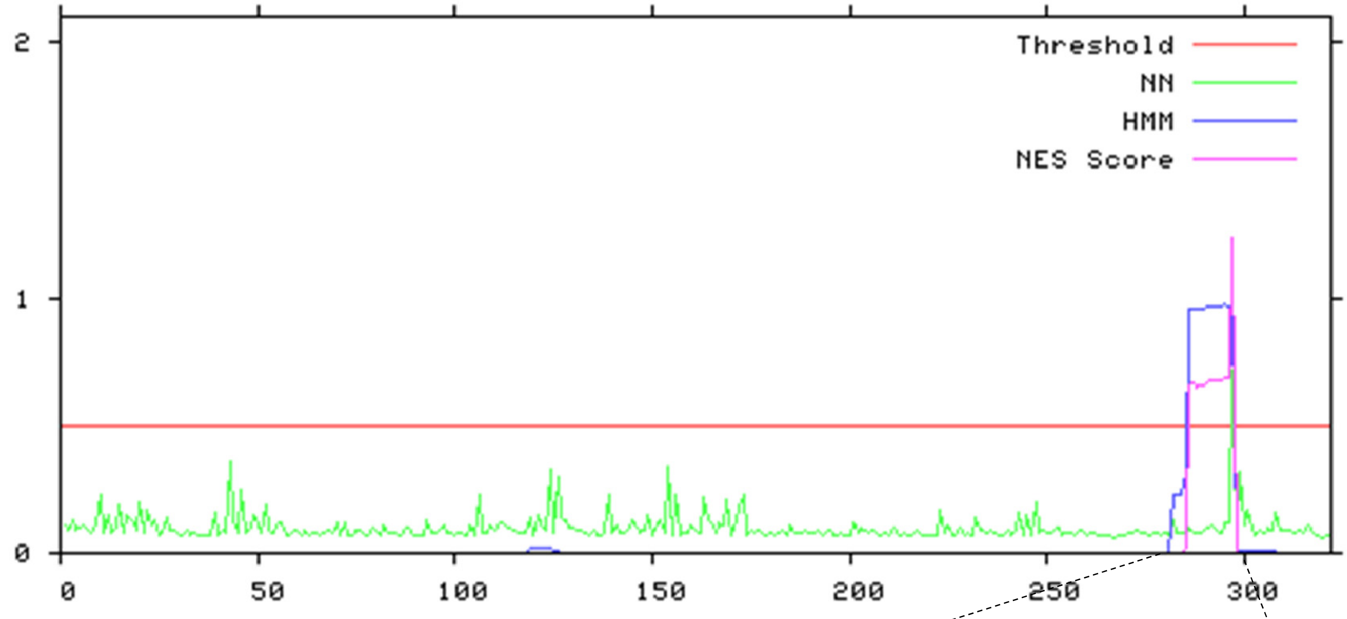


NetNES optimized prediction: LRPF**V**PEQE**LL**RS**L**R**L**

Hidden Markov Model (HMM) algorithm prediction: **L**RP**FV**PEQE**LL**RS**L**R**L**

Supplementary Figure S2. NetNES prediction for NES in αTAT1 identifying the putative NES, the predicted NES from NetNES optimized algorithm (pink) and that from Hidden Markov Model (blue) are shown with the hydrophobic residues indicated in bold.

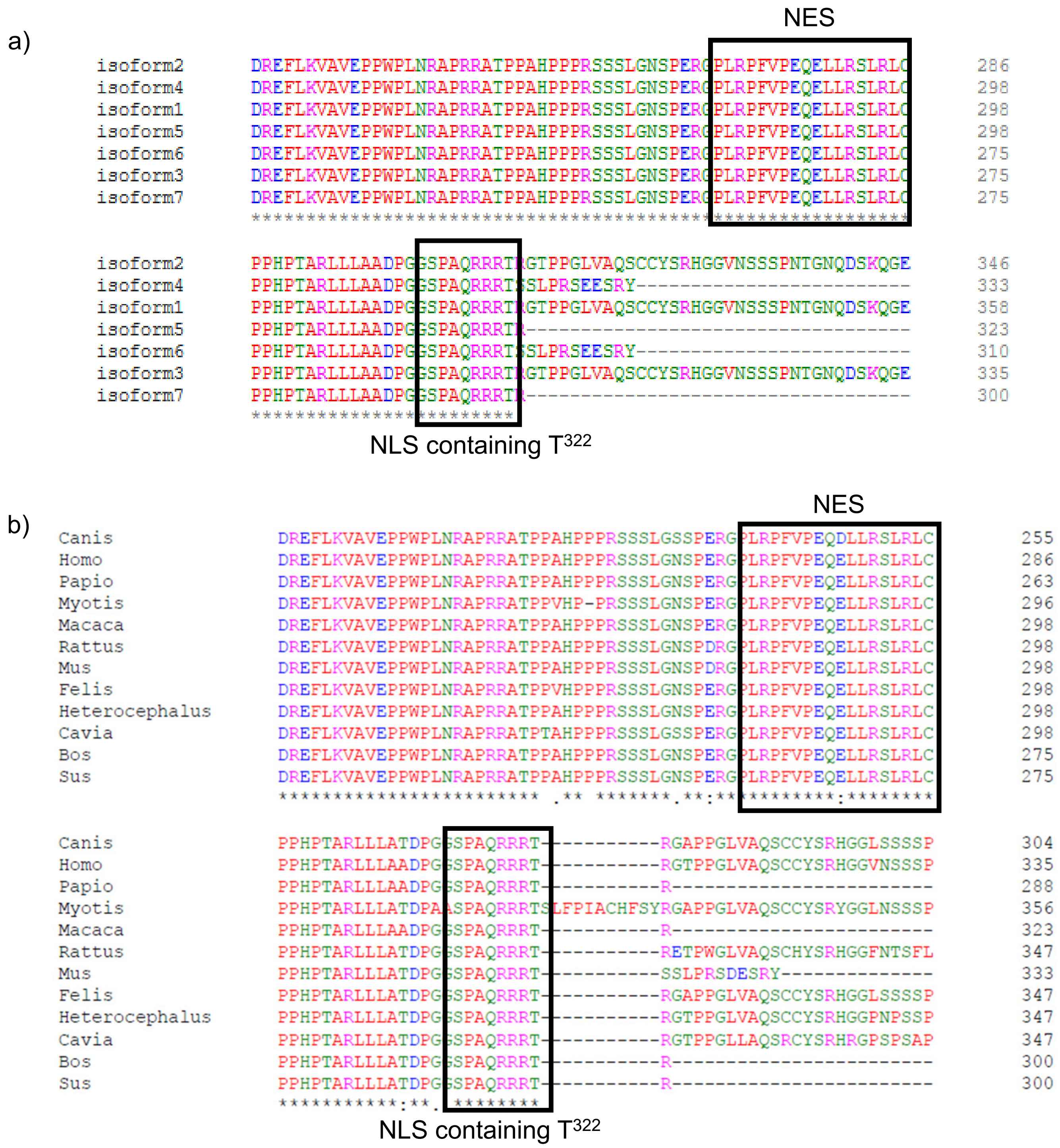


Supplementary Figure S3. The putative NES and NLS in α-TAT1 are evolutionarily conserved. a) Alignment of human α-TAT1 isoforms, b) alignment of different mammalian α-TAT1 showing the C-terminal region including the putative NES and NLS (enclosed in boxes), genus of the mammalian α-TAT1 are shown. Alignments were performed using ClustalW (https://www.ebi.ac.uk/Tools/msa/clustalo/)

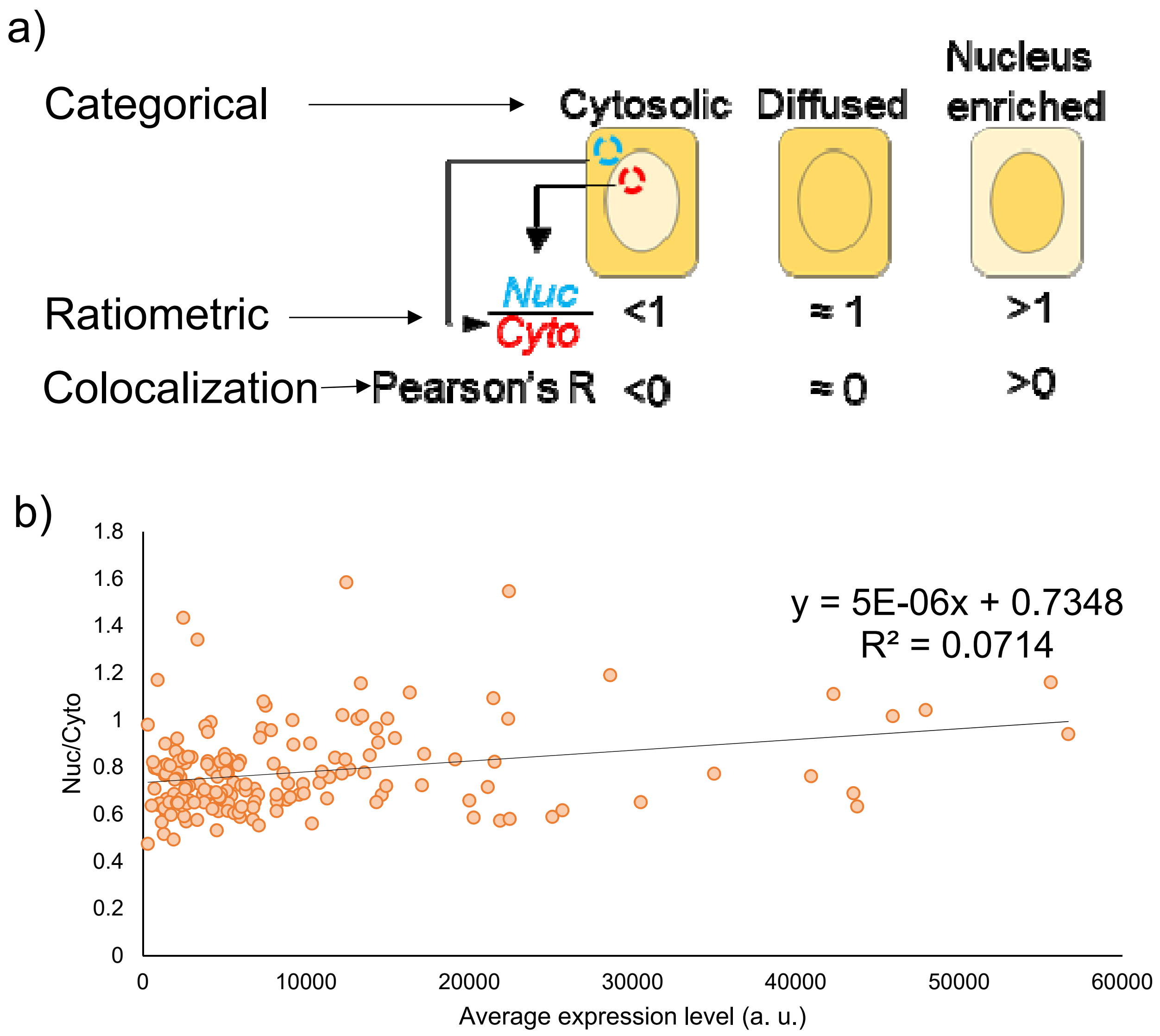


Supplementary Figure S4. a) Strategies to quantify intracellular localization of mVenus-α-TAT1, b) Changes in Nuc/Cyto intensity ratio of mVenus-αTAT1 with respect to expression levels in transiently transfected HeLa cells.

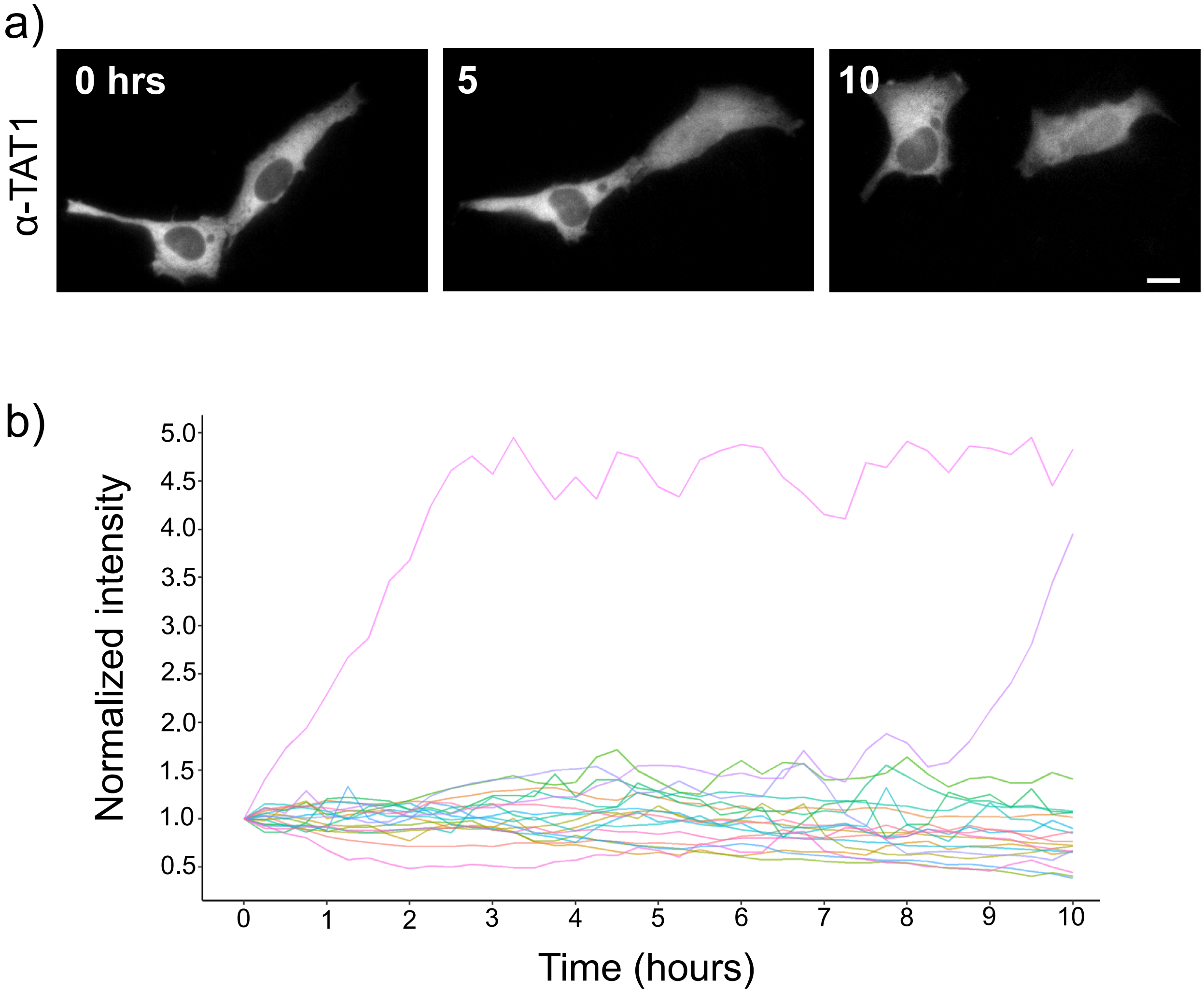


Supplementary Figure S5. α-TAT1 intracellular localization is dynamic in nature. a) Spontaneous temporal changes in spatial distribution of mVenus-α-TAT1, scale bar = 10 μm b) temporal changes in nuclear fluorescence intensity of mVenus-α-TAT1 in HeLa cells, each line indicates normalized nuclear intensity of mVenus- α-TAT1 in a single cell, n = 20 cells

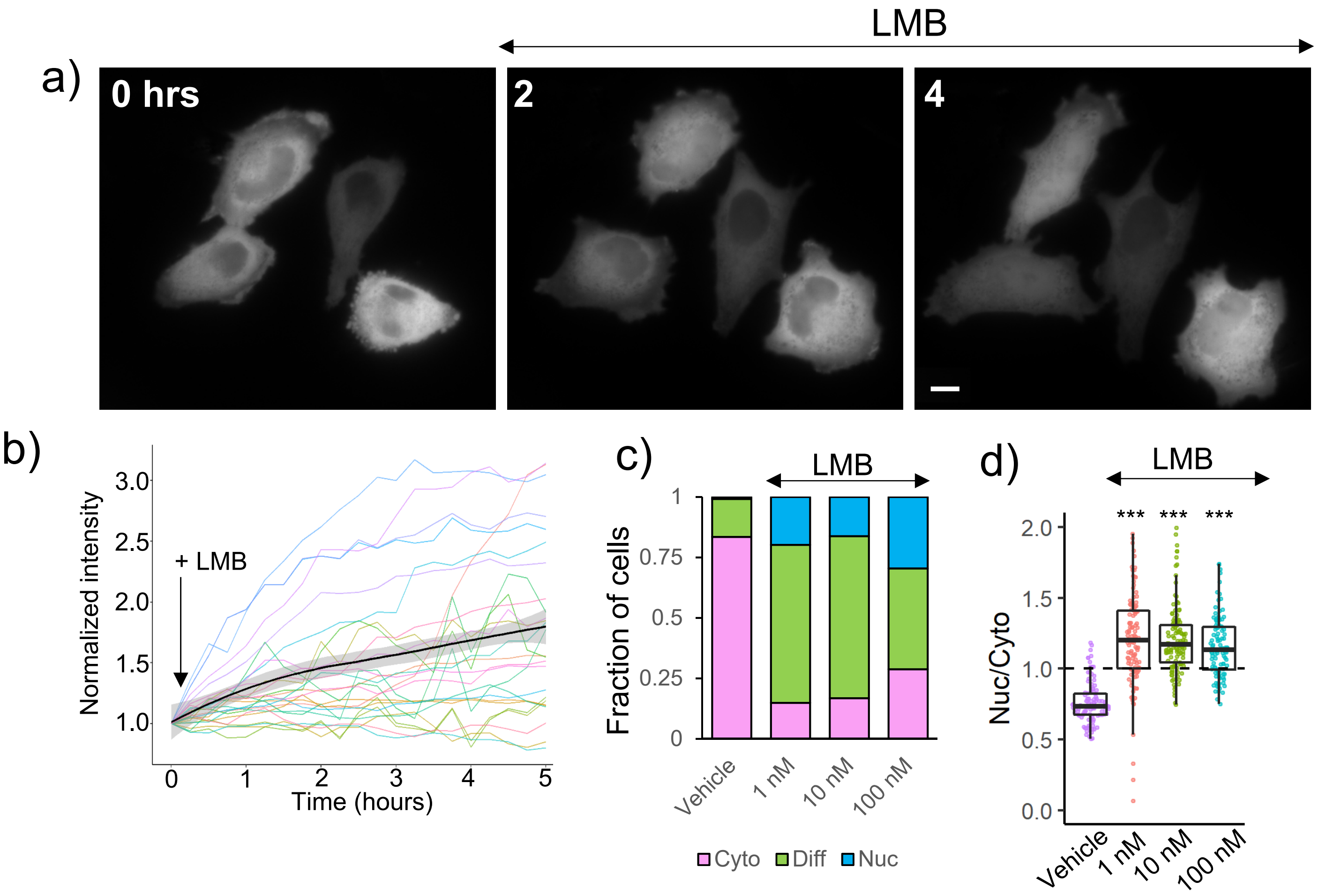


Supplementary Figure S6. α-TAT1 undergoes CRM1 dependent nuclear export. a), b) Temporal changes in mVenus-α-TAT1 localization on 100 nM LMB treatment after LMB addition at time 0, for b), each line indicates normalized nuclear intensity of mVenus- α-TAT1 in a single cell, black line shows the mean and gray line indicate the 95% C.I., n = 26 cells , c) categorical analysis ( vehicle: 274, LMB 1 nM: 329, LMB 10 nM: 291, LMB 100 nM: 295 cells) and d) ratiometric analysis of mVenus-α-TAT1 localization on vehicle (103 cells) or LMB treatment at 1 nM (121 cells), 10 nM (122 cells) and 100 nM (103 cells). Scale bar = 10μm. P-value: *** <0.001 or as shown, Student's *t*-test.

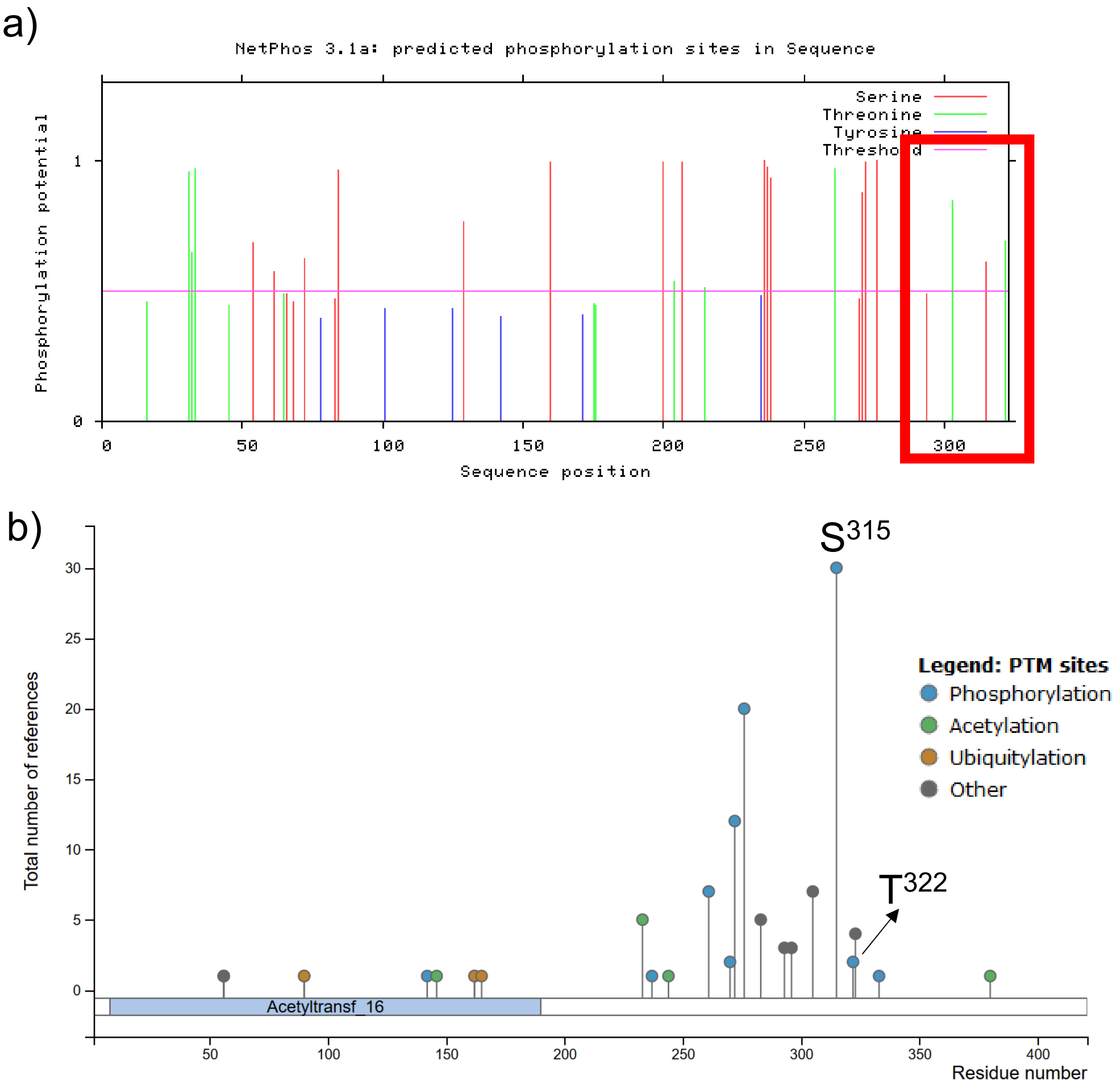


Supplementary Figure S7. a) Putative phosphosites in α-TAT1 predicted by NetPhos, those in F285-R323 are highlighted, b) currently reported post-translational modifications in α-TAT1 curated by PhosphoSitePlus®, www.phosphosite.org.

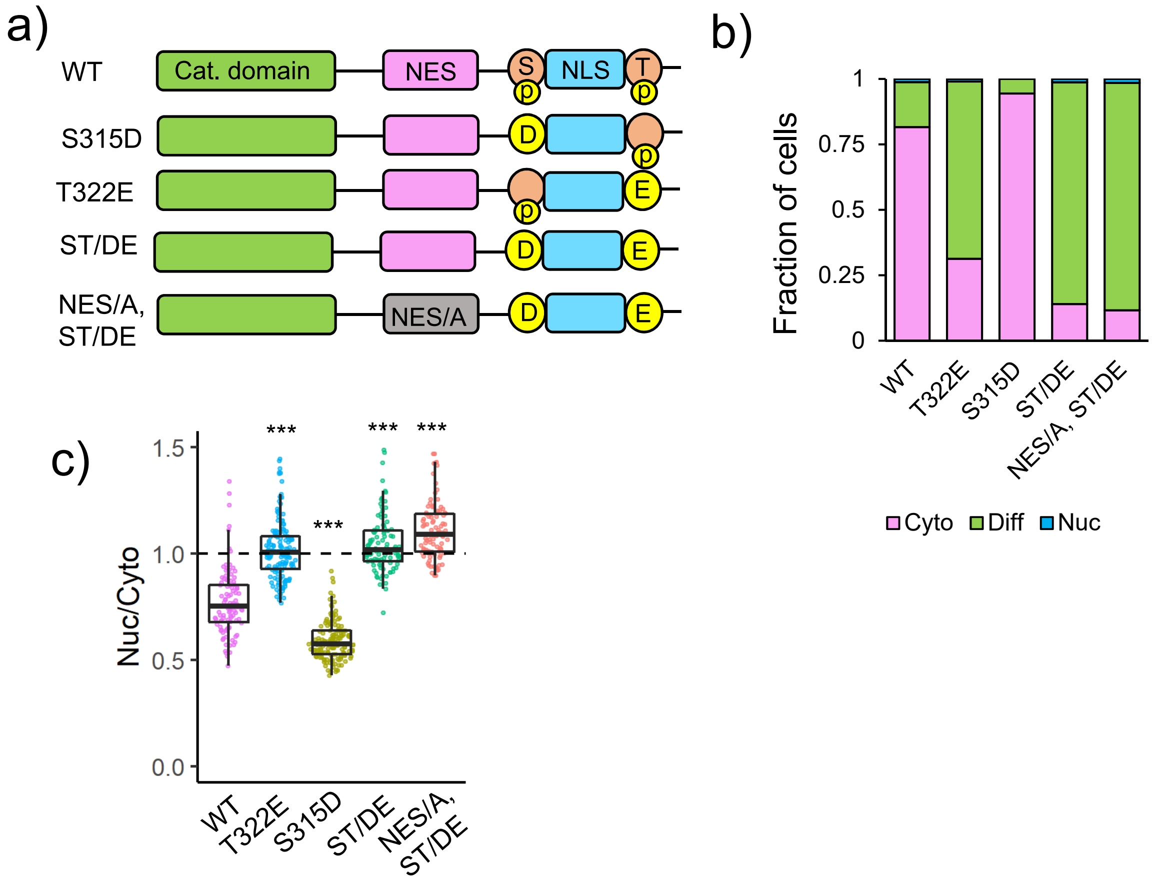


Supplementary Figure S8. α-TAT1 has a C-terminal phospho-inhibited NLS. a) Cartoon showing α-TAT1 NES and NLS flanked by potential phosphosites and mutants, b) categorical (WT:283, T322E: 248, S315D: 227, ST/DE: 234, NES/A-ST/DE: 238 cells) and c) ratiometric (WT:104, T322E: 148, S315D: 154, ST/DE: 103, NES/A-ST/DE: 98 cells) analyses of intracellular localization of mVenus-α-TAT1 mutants. ****P*<0.001, Student's *t*-test.

a)

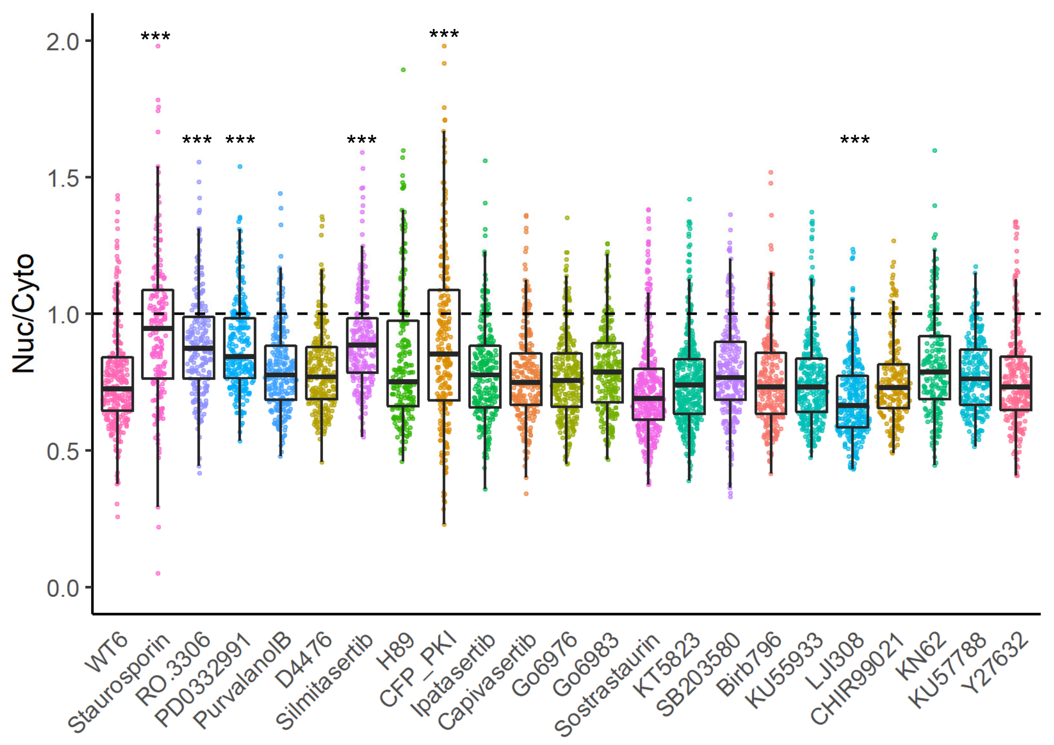


b)

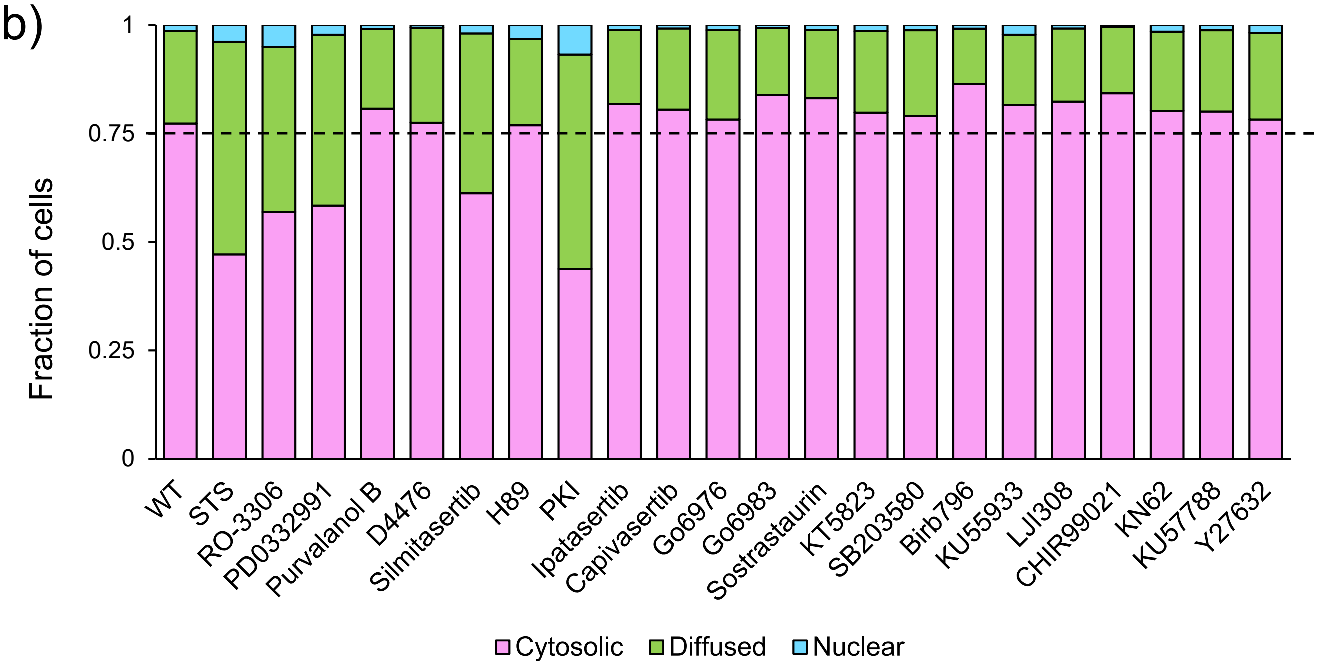


Supplementary Figure S9. a) Ratiometric and b) categorical analysis of α-TAT1 localization on treatment with kinase inhibitors. WT: 1032, STS: 157, R0-3306: 796, PD0332991: 543, Purvalanol B: 992, D4476: 689, 418, H89: 251, PKI: 354, Ipatasertib: 546, Capivasertib: 385, Go6976: 712, Go6983: 606, Sostrastaurin: 605, KT5823: 570, SB203580: 505, Birb796: 991, KU55933: 815, LJI308: 522, CHIR99021: 484, KN62: 531, KU57788: 949, Y27632: 340 cells. ****P*<0.001, Student’s *t*-test.

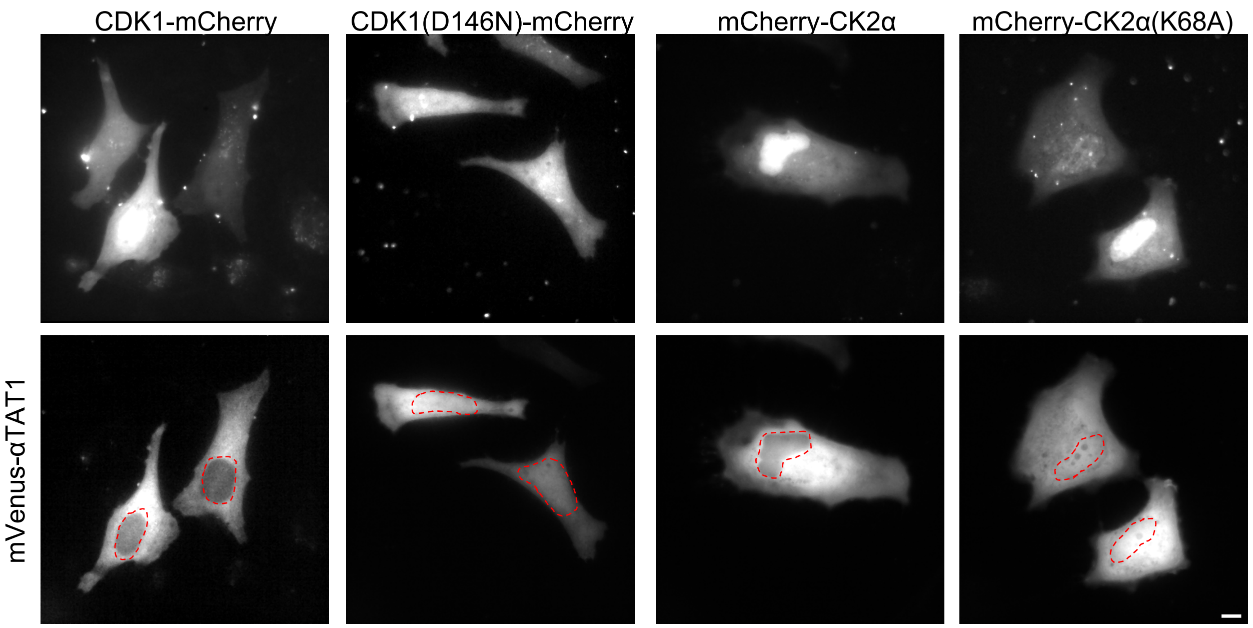


Supplementary Figure S10. Localization of mVenus-αTAT1 on co-expression with Cdk1-mCherry, Cdk1(D146N), mCherry-CK2α and mCherry-CK2α(K68A). Hoescht staining was performed to identify nuclei. Scale bar= 10 μm

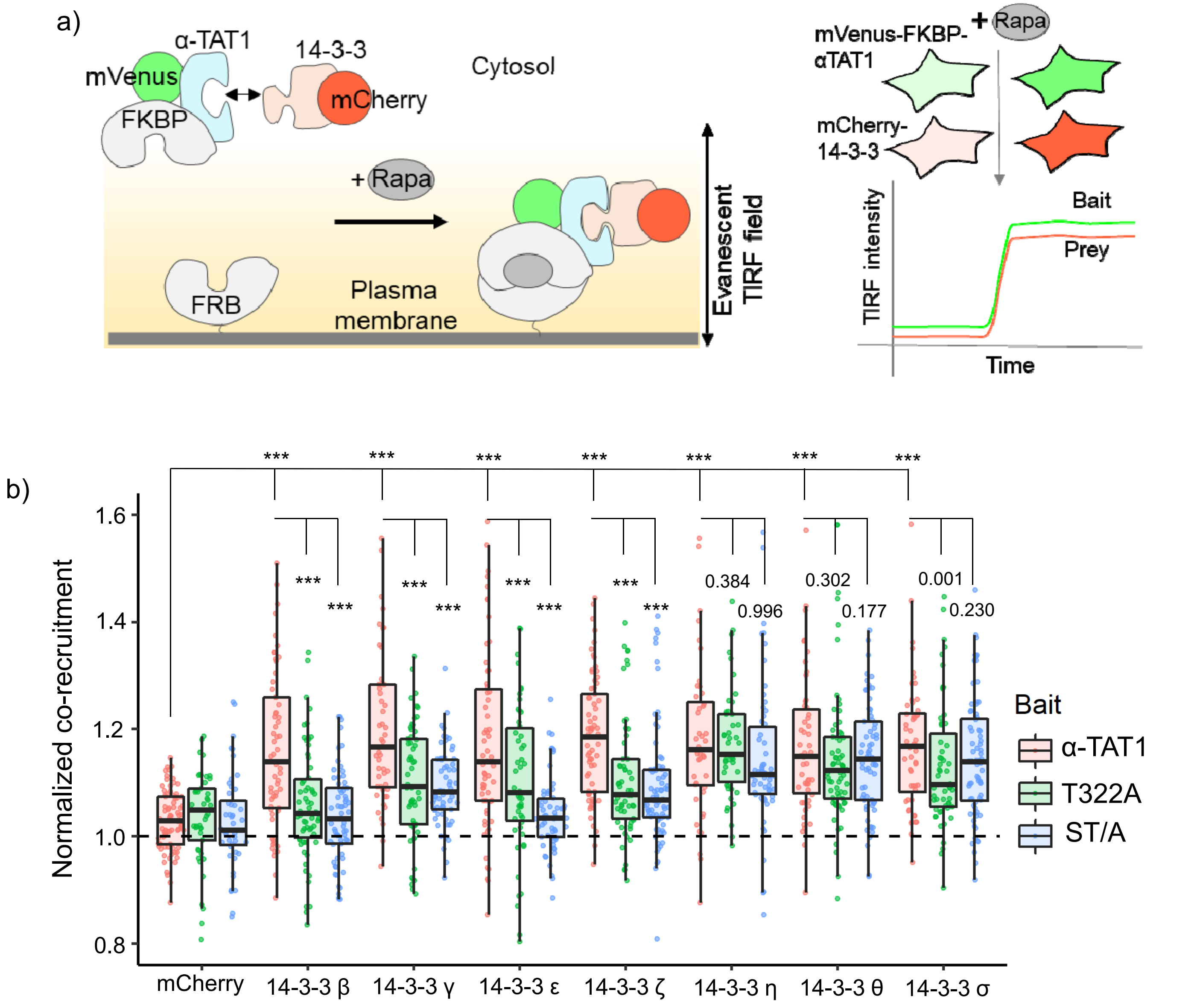


Supplementary Figure S11. a) Schema of chemically-inducible co-recruitment assay. Rapamycin-induced interaction between FKBP and FRB was used to assess the extent of binding of prey candidates against a bait. (Left) FKBP is fused to the prey along with a fluorescent protein, while FRB is tethered to an inner leaflet of plasma membrane. Upon rapamycin addition, FKBP binds to FRB which brings the bait (mVenus-FKBP-α-TAT1) and the prey capable of binding (mCherry-14-3-3) to the plasma membrane. (Right) Recruitment of the bait and the prey to the plasma membrane can be sensitively detected by TIRF microscopy as an increased fluorescence signal. (b) Normalized co-recruitment levels by indicated baits of mCherry (α-TAT1: 79, T322A: 45, ST/A: 49), mCherry-14-3-3β (α-TAT1: 64, T322A: 63, ST/A: 66), mCherry-14-3-3γ (α-TAT1: 50, T322A: 57, ST/A: 64), mCherry-14-3-3ε (α-TAT1: 65, T322A: 52, ST/A: 58), mCherry-14-3-3ζ (α-TAT1: 70, T322A: 58, ST/A: 79), mCherry-14-3-3η (α-TAT1: 48, T322A: 50, ST/A: 58), mCherry-14-3-3θ (α-TAT1: 53, T322A: 68, ST/A: 53), mCherry-14-3-3σ (α-TAT1: 53, T322A: 56, ST/A: 73), P-value: *** <0.001 or as shown, Student's *t*-test.

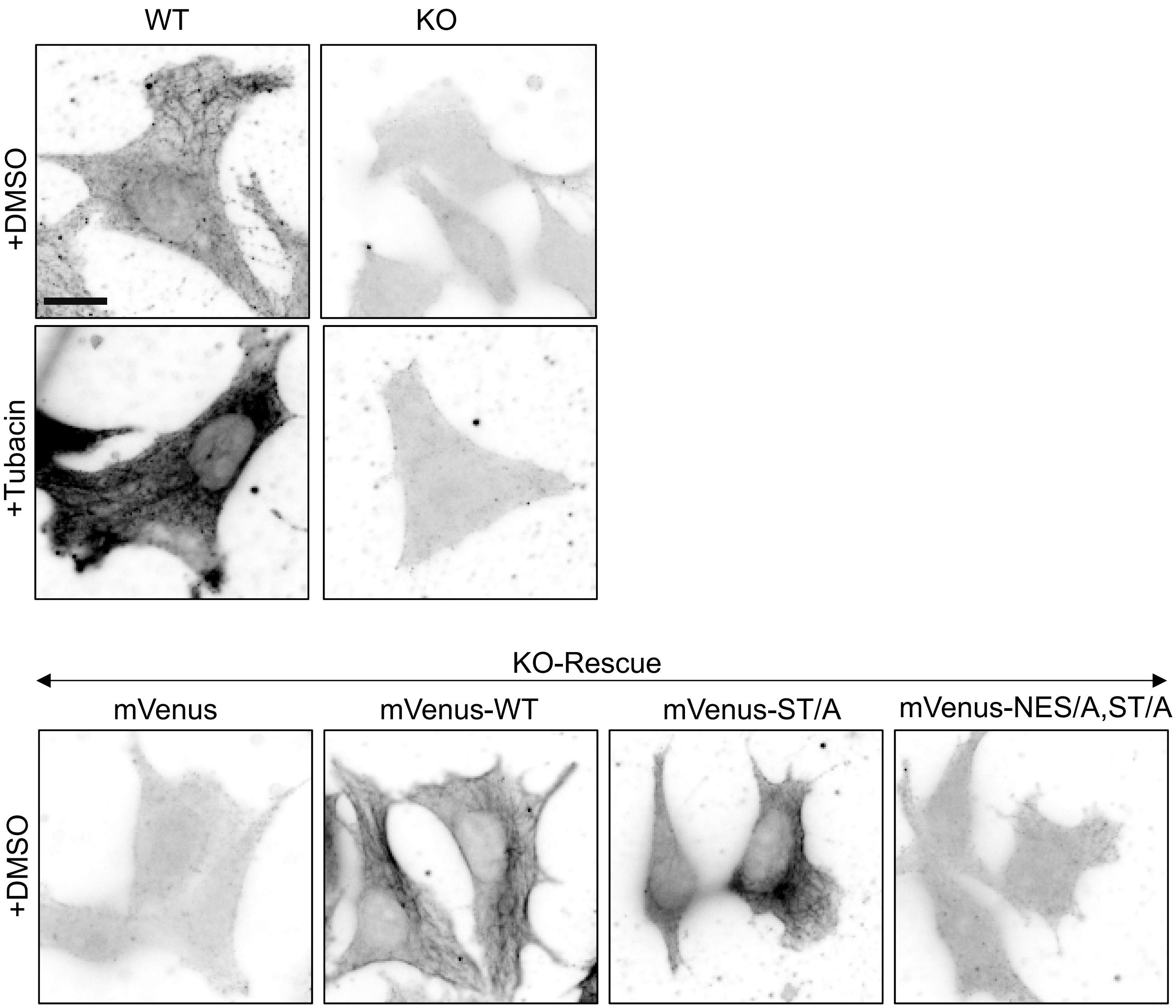


Supplementary Figure S12. MT acetylation levels in WT and α-TAT1 KO MEFs with DMSO(vehicle) or Tubacin, and KO MEFs expressing mVenus, mVenus-α-TAT1, mVenus-α-TAT1(ST/A) and mVenus-α-TAT1(NES/A, ST/A) treated with DMSO. Scale bar: 10 μm.

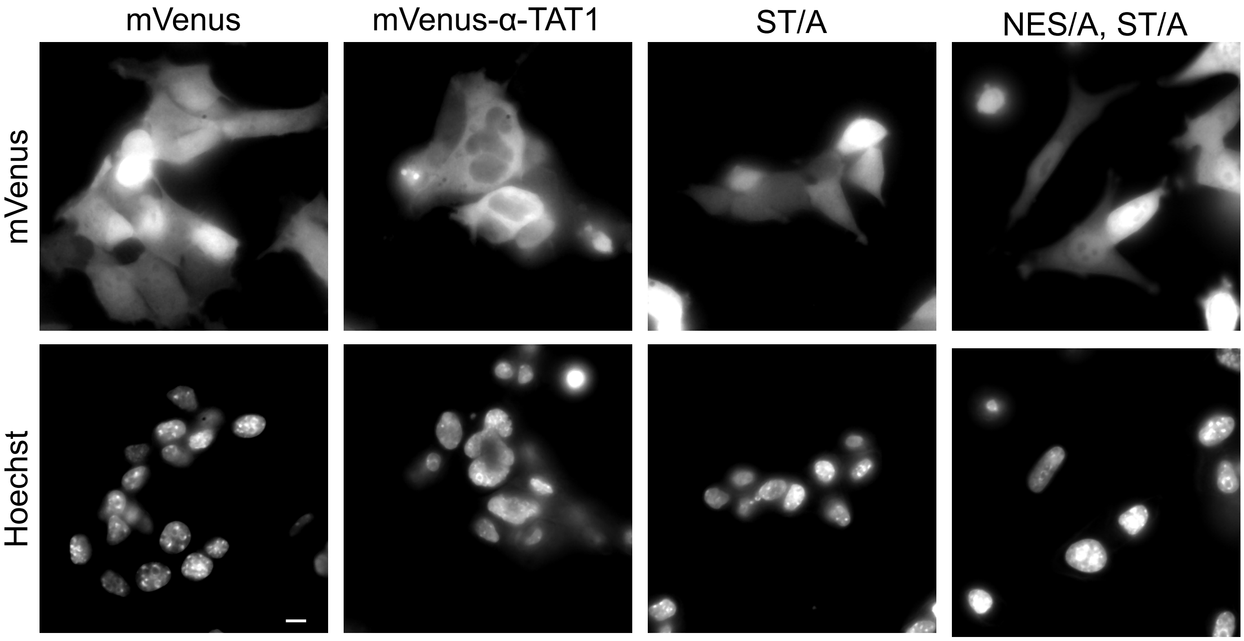


Supplementary Figure S13. Localization of mVenus, mVenus-αTAT1, mVenus-αTAT1(ST/A) and mVenus-αTAT1(NES/A, ST/A) stably expressed in αTAT1 knock-out MEFs. Hoechst stain was used to identify nuclei. Scale bar= 10 µm

a)

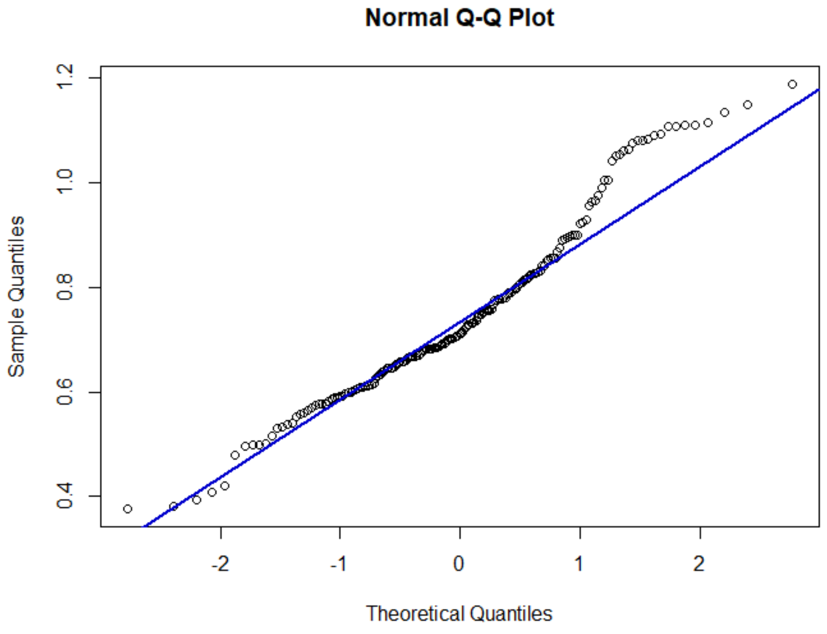


b)

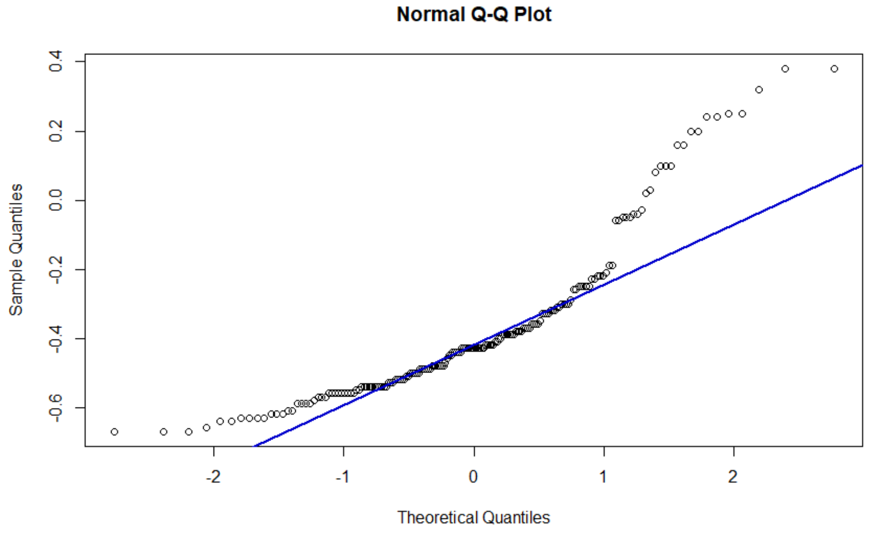


Supplementary Figure S14. Normal probability plot of a) nuclear/cytosolic ratio of mVenus-α-TAT1 and b) Pearson's R value between mVenus-α-TAT1 and Histone-2B-mCherry expressed in HeLa cells.

| Drug | Conc. Used | target kinase | Incubation time | p value (t-test) |
|---|---|---|---|---|
| STS | 100 nM | Pan kinase | 4 hours | 1.20187E-11 |
| RO3306 | 10 µM | Cdk1, Cdk2 | 4 hours | 1.87845E-10 |
| PD0332991 | 1 µM | Cdk4, Cdk6 | 4 hours | 1.29985E-10 |
| Purvalanol B | 1 µM | Cdk1, Cdk2, Cdk5 | 4 hours | 0.086698844 |
| D4476 | 1 µM | CK1 | 4 hours | 0.244864719 |
| Silmitasertib | 10 µM | CK2 | 4 hours | 7.34743E-12 |
| H89 | 10 µM | PKA | 4 hours | 0.001868139 |
| PKI | NA | PKA | Overnight | 8.76993E-08 |
| Ipatasertib | 10 µM | PKB (Akt) | 4 hours | 0.026614985 |
| Capivasertib | 1 µM | PKB (Akt) | 4 hours | 0.928873535 |
| Go6976 | 250 nM | PKC | 4 hours | 0.900843024 |
| Go6983 | 250 nM | PKC | 4 hours | 0.068144954 |
| Sostrastaurin | 10 µM | PKC | 4 hours | 0.008916116 |
| KT5823 | 5 µM | PKG | 4 hours | 0.581314971 |
| SB203580 | 10 µM | p38 MAPK | 4 hours | 0.007945499 |
| Birb796 | 10 µM | p38 MAPK | 4 hours | 0.768174955 |
| KU55933 | 10 µM | ATM | 4 hours | 0.26994365 |
| LJI308 | 10 µM | RSK | 4 hours | 9.62544E-07 |
| CHIR99021 | 1 µM | GSK3 | 4 hours | 0.309440886 |
| KN62 | 10 µM | CaMK-II, P2RX7 | 4 hours | 0.021582087 |
| KU57788 | 1 µM | DNA-PK | 4 hours | 0.385979985 |
| Y27632 | 10 µM | ROCK | 4 hours | 0.375473038 |

Supplementary Table T1. Details of kinase inhibitor screening assay.
